# New genetic tools enable dissection of a global stress response in the early-branching species *Fusobacterium nucleatum*

**DOI:** 10.1101/2022.07.29.501972

**Authors:** Falk Ponath, Yan Zhu, Valentina Cosi, Jörg Vogel

## Abstract

*Fusobacterium nucleatum*, long known as a common oral microbe, has recently garnered attention for its ability to colonize tissues and tumors elsewhere in the human body. Clinical and epidemiological research has now firmly established *F. nucleatum* as an oncomicrobe associated with several major cancer types. However, with the current research focus on host associations, little is known about gene regulation in *F. nucleatum* itself, including global stress response pathways that typically ensure the survival of bacteria outside their primary niche. This is due to the phylogenetic distance of Fusobacteriota to most model bacteria, their limited genetic tractability, and paucity of known gene functions. Here, we characterize a global transcriptional stress response network governed by the extracytoplasmic function sigma factor, σ^E^. To this aim, we developed several new genetic tools for this anaerobic bacterium, including four different fluorescent marker proteins, inducible gene expression, scarless gene deletion, and transcriptional and translational reporter systems. Using these tools, we identified a σ^E^ response partly reminiscent of phylogenetically distant Proteobacteria but induced by exposure to oxygen. Although *F. nucleatum* lacks canonical RNA chaperones such as Hfq, we uncovered conservation of the non-coding arm of the σ^E^ response in form of the non-coding RNA FoxI. This regulatory small RNA (sRNA) acts as an mRNA repressor of several membrane proteins, thereby supporting the function of σ^E^. In addition to the characterization of a global stress response in *F. nucleatum*, the genetic tools developed here will enable further discoveries and dissection of regulatory networks in this early-branching bacterium.

**SIGNIFICANCE STATEMENT:** *Fusobacterium nucleatum* is an abundant member of the oral microbiome that can spread throughout the body and colonize secondary sites, including cancer tissues where it promotes tumor progression. Understanding how *F. nucleatum* is able to adapt to this new environment might open new therapeutic opportunities, but we currently lack basic molecular knowledge of gene regulation in this phylogenetically distinct bacterium. We developed much-needed genetic tools for use in *F. nucleatum* and with their aid uncovered a stress response mediated by the transcriptional activator σ^E^ and an associated small RNA. Our findings in an early-branching bacterium reveal surprising parallels to and differences from the σ^E^ response in well-characterized model bacteria and provide a framework that will accelerate research into the understudied phylum Fusobacteriota.

## INTRODUCTION

*Fusobacterium nucleatum* is an abundant member of the oral microbiome, a complex microbial community consisting of over 700 species (1). In the oral cavity, *F. nucleatum* functions as a bridging organism between colonizers of dental plaque (2) and is crucial for the maintenance of this biofilm (3). While generally considered a mutualist, *F. nucleatum* is also implicated in periodontitis and occurs in abscesses in various body sites (4). Most importantly, *F. nucleatum* has recently garnered broad attention for its association with several different types of human tumors. The bacterium is highly abundant in tissues of colorectal, breast, esophageal and pancreatic cancer (5–11), and colonization with *F. nucleatum* is associated with enhanced tumor growth, metastasis, and resistance to chemotherapy (9–11). These tissues represent new and adverse environments very different from *F. nucleatum*’s primary niche in the oral cavity.

The occurrence of *F. nucleatum* at these extraoral sites implies that this bacterium can sense and adapt to changes in its environment. To date, however, only two regulatory circuits have been described in *F. nucleatum*: The two-component systems (TCS) CarRS and ModRS, which control interspecies co-aggregation and resistance to hydrogen peroxide, respectively. Both also play a role in bacterial virulence (12, 13). Factors that govern global stress responses are unknown. Decoding molecular principles of gene regulation in *F. nucleatum* has generally been difficult for two reasons. First, the phylum Fusobacteriota is phylogenetically remote from all model bacteria (14) (Fig. 1A), which hampers knowledge transfer by sequence comparison. Second, functional genetics in this obligate anaerobic, Gram-negative bacterium is in its infancy (2), being limited to two recently-introduced systems for scarless genomic deletion (15, 16) and to plasmid-based overexpression of a gene of interest (17). Therefore, new genetic tools are much needed to systematically identify and characterize regulatory pathways that ensure viability of *F. nucleatum* under stress conditions. In this work, we expand the fusobacterial genetic tool-kit and use it to dissect a global stress response composed of the extracytoplasmic function (ECF) σ factor, σ^E^, and an associated regulatory small RNA (sRNA).

**Figure 1:**
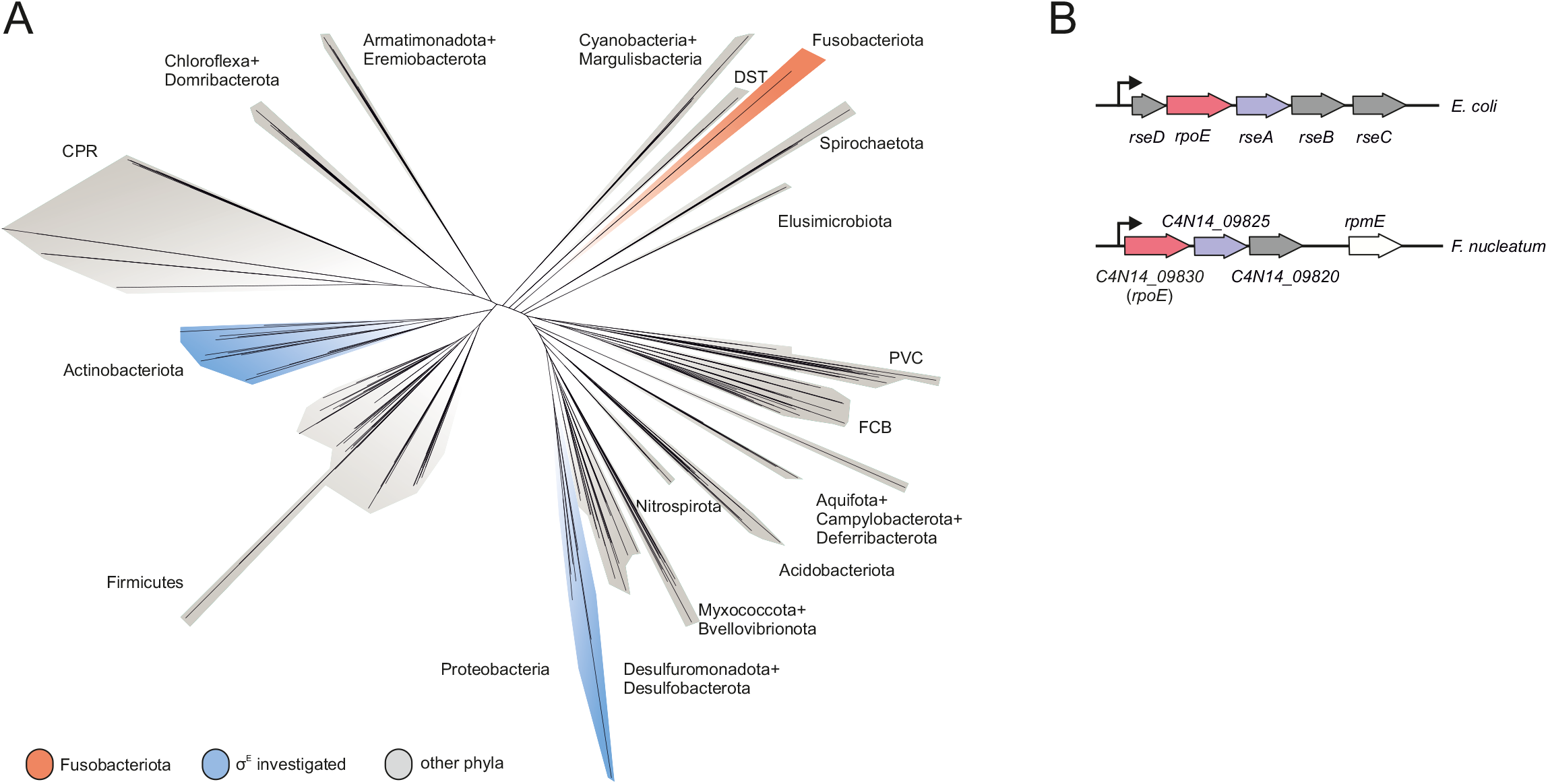
Phylogenetic positioning of Fusobacteriota and comparison of the ECF locus. (A) A phylogenetic tree of 265 bacterial species based on the alignments provided by *Coleman et al.* (2021) (14). (B) Schematic representation of the *rpoE* operon in *E. coli* and *F. nucleatum*. *rpoE* genes in red; the anti-sigma factor *rseA* and its putative homologue in *F. nucleatum* in purple; the remaining gene(s) in the respective *rpoE* operons in grey.

ECFs present a fundamental signal transduction mechanism whereby bacteria monitor their environment (18). Specifically, ECFs are usually involved in regulating the integrity of the bacterial envelope (19), which represent the first line of defense in Gram-negative bacteria (20). ECFs across different phyla are activated by a variety of signaling stimuli, including broad stresses, such as osmotic stress, heat shock, oxidative stress (20), but also more specific stressors, such as singlet oxygen produced by photosynthesis (21) or lysozyme (22). Nevertheless, ECFs share similar core features: (i) they are auto-regulatory, (ii) an anti-sigma factor that keeps the ECF in an inactive state is encoded in the same operon, (iii) upon sensing of the specific signal the anti-sigma factor is inactivated either through proteolytic degradation (23), conformational changes (24, 25) or sequestration by a third protein (26). More rarely, ECFs can also be activated through phosphorylation (27) or via a TCS (28).

The σ^E^ response has been particularly well-studied since its discovery in *Escherichia coli* more than three decades ago (29). Upon envelope stress, proteolytic cleavage and degradation of the anti-sigma factor RseA releases σ^E^ from the inner membrane (IM) into the cytosol, where σ^E^ activates the transcription of >100 genes (30, 31). The σ^E^ regulon is functionally similar across different bacterial species and includes genes involved in DNA damage repair, LPS biogenesis and outer membrane (OM) homeostasis (31–35).

Importantly, the σ^E^ regulon also includes several sRNAs genes, which together constitute the ‘non-coding arm’ of the response (36). Work in *E. coli*, *Salmonella* and *Vibrio* species has established that these σ^E^-controlled sRNAs post-transcriptionally repress mRNAs of diverse envelope proteins, including many major outer membrane proteins (OMPs) (37–45). Endowing the transcriptional activator σ^E^ with a repressor function, these sRNAs act synergistically to ensure envelope integrity. In all species investigated thus far, these sRNAs work in conjunction with the general RNA chaperone Hfq, which aids base pairing between the sRNAs and their target mRNAs. In fact, chronic activation of the σ^E^ response as a result of perturbed envelope homeostasis is a conserved characteristic of *hfq* deletion strains among these species (46–49).

The envelope composition of *F. nucleatum* is largely unknown, and there has been conflicting evidence with respect to a potential σ^E^ stress response. First, genome annotation of *F. nucleatum* (50) predicted a putative *rpoE* gene, which encodes σ^E^ in *E. coli* (Fig. 1B). However, a recent comprehensive phylogenetic analysis placed the putative *F. nucleatum* σ^E^ in a functionally different ECF group from the *E. coli* protein (51). Second, our recent RNA-seq study in *F. nucleatum* discovered a previously unknown large suite of sRNAs. Preliminary analysis identified one of these sRNAs, FoxI, as a post-transcriptional repressor of an abundant OMP (17). However, FoxI was induced by molecular oxygen, a condition which seems unrelated to envelope stress and untypical of a σ^E^ response. More importantly, *F. nucleatum* lacks a gene coding for Hfq or any other known sRNA chaperone.

Here, to experimentally resolve these seeming inconsistencies, we developed several much-needed systems to characterize fusobacterial gene regulation: fluorescent marker proteins, transcriptional and translational reporters, an inducible gene expression system and a gene deletion system that is not reliant on a specific strain background. Application of these tools allowed us to define the σ^E^ regulon of *F. nucleatum,* revealing a surprising conservation of its overall architecture in this early-branching species. This general conservation includes a non-coding arm of the σ^E^ response provided by the sRNA FoxI, which we show acts as a negative post-transcriptional regulator of several envelope proteins. Intriguingly, the fusobacterial ECF is activated by oxygen rather than sources of envelope or oxidative stress. Our results provide the first functional evidence for a global stress response composed of a sigma factor and an associated small RNA in an early-branching bacterium, and a new experimental framework to dissect regulatory networks in the understudied phylum Fusobacteriota.

## RESULTS

### Expanding the genetic toolkit for F. nucleatum

To facilitate dissection of gene regulatory networks in *F. nucleatum*, we created five plasmid-based tools: constitutively expressed fluorescent marker proteins, a transcriptional and a translational reporter system, an inducible gene expression system, and a system for markerless genomic deletion (Fig. 2A). These systems are derived from our recently developed plasmid pEcoFus for gene overexpression in *F. nucleatum* (17). We initially reduced the overall size of pEcoFus, generating the plasmid pVoPo-00 (see Methods for details). Next, we inserted an expression cassette for one of four different codon-optimized fluorescent proteins: mCherry, GFP, mScarlet-I and mNeonGreen. These constructs allow easy visualization of *F. nucleatum* using fluorescence imaging (Fig. 2B). pVoPo-mNeonGreen uses an expression construct with a weaker promoter, therefore the fluorescence signal is lower (see Material and Methods). This adjustment was necessary because we were unable to express the original mNeonGreen construct in *E. coli* during the cloning procedure, likely due to overexpression toxicity. The pVoPo-mNeonGreen construct also demonstrates that marker gene expression can be adjusted and thereby adapted to the conditions needed. In addition to their excitation and emission spectra, the four fluorescence proteins also differ in photostability and maturation time. This will allow researchers to choose the optimal vector for their specific experimental needs.

**Figure 2:**
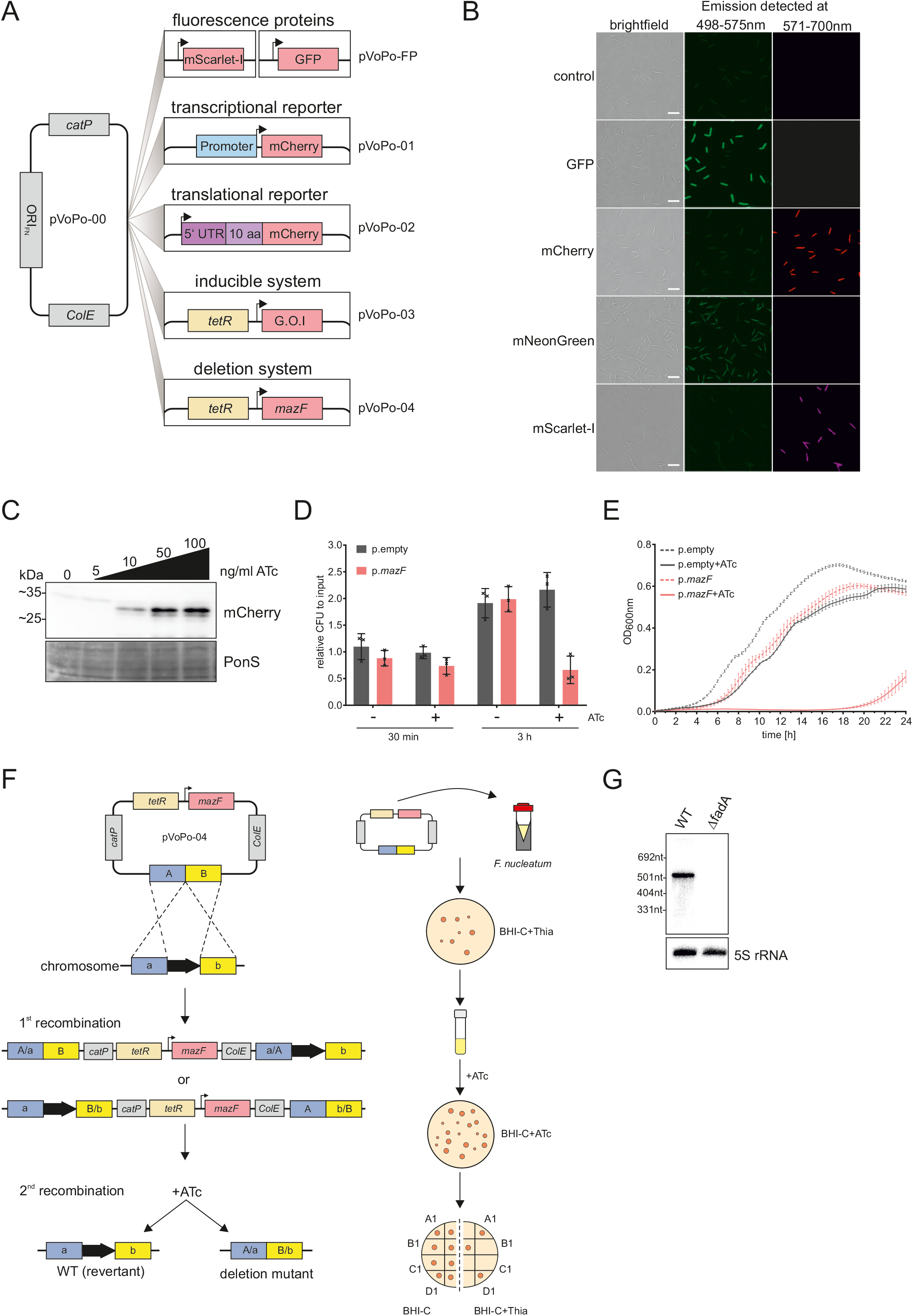
A genetic toolbox for *F. nucleatum*. (A) Overview of the plasmid-based genetic tools developed for *F. nucleatum* based on the vector pVoPo-00. FP, fluorescent protein (GFP, mCherry, mNeonGreen, mScarlet-I); G.O.I, gene of interest; catP, chloramphenicol resistance cassette; ColE, replication of origin; ORI_FN_, origin of replication for *F. nucleatum*; tetR, tetracycline repressor. (B) Representative images of *F. nucleatum* carrying pVoPo-FP expressing different fluorescent proteins. The cells were PFA-fixated prior to overnight maturation at 4 °C. The emission was detected at the indicated wavelength. Scale bar, 5 µM (C) Western blot analysis of lysates of *F. nucleatum* carrying a plasmid with *mCherry* introduced in the inducible system (pVoPo-03) after 30 min exposure to different ATc concentrations. mCherry runs as a duplet likely representing the full-length protein as well as a truncated version arising from an internal translational start site (53). ATc, anhydrotetracycline. Ponceau S (PonS) staining is shown as loading control. (D) Quantification of colony-forming units (CFU) for *F. nucleatum* carrying the empty vector control pVoPo-03 (p.empty) or the pVoPo-03-*mazF* plasmid (p.*mazF*). The bacteria were grown to mid-exponential phase and treated with 100 ng ml^-1^ ATc to induce *mazF* expression. Serial dilutions of the samples were plated after 0 min (input), 30 min and 3 h. Untreated samples were used as control. Data are presented as the average and standard deviation for three biological replicates relative to the input CFUs. (E) Growth curves for *F. nucleatum* carrying either p. empty or p.*mazF* in the presence or absence of 100 ng ml^-1^ ATc. No selection pressure for plasmid maintenance was included. Displayed is the average of three biological replicates with standard deviation. (F) Schematic representation of allelic exchange (left side) and experimental workflow (right side) using the pVoPo-04 system to generate unmarked deletion strains. A/B, up- and downstream homology regions. (G) Northern blot detection of *fadA* using total RNA samples extracted from *F. nucleatum* wild-type (WT) or a Δ*fadA* strain generated via the deletion system pVoPo-04. 5S rRNA served as loading control.

On the basis of pVoPo-mCherry, we generated the transcriptional reporter plasmid pVoPo-01 to determine the activity of promoter regions of interest placed upstream of mCherry (Fig. 2A). For translational reporters (plasmid pVoPo-02), the 5’ UTR and first 10 amino acids (aa) of a specific target gene are fused to the second codon of mCherry and expressed under a constitutive promoter (Fig. 2A).

Next, we adapted the widely used tetracycline-inducible gene expression system (52) for use in *F. nucleatum*. To this end, we replaced the constitutive promoter of pVoPo-00 with a synthetic tetracycline-responsive promoter and added a TetR repressor gene, thus creating plasmid pVoPo-03. For proof of concept, we showed that expression of mCherry from this plasmid can be tightly controlled with the non-bacteriostatic tetracycline derivative, anhydrotetracycline (ATc). Addition of different concentrations of ATc to cultures of *F. nucleatum* led to a dose-dependent and robust expression of mCherry (Fig. 2C), whereas no signal was detected in the absence of ATc. Of note, in the western blots, mCherry runs as a doublet, likely representing full-length protein and a truncated version expressed from an internal translational start site (53).

Gene deletion represents another important tool in the arsenal to study gene regulation in bacteria. The currently available gene disruption (54, 55) and deletion (15) tools for *F. nucleatum* use suicide vectors. However, these vectors require either constant selection pressure (56, 57) or a *galK(T)* gene deletion background for efficient counter-selection (15, 16), which limits their application for long-term or complex experiments, such as animal studies. Taking advantage of the inducible plasmid pVoPo-03, we evaluated the MazF toxin as a potential counter-selection marker, since heterologous expression of this endonuclease had been shown to be toxic in several unrelated bacteria (58–60). Similarly, in *F. nucleatum*, we observed a 40% reduction in viable bacteria 3 hours (h) after *mazF* induction (Fig. 2D). Induced expression of MazF caused a drastic growth delay for ∼16h, before growth resumed (Fig. 2E). Importantly, no recovery was observed when this experiment was performed under conditions that select for plasmid retention (Supplementary Fig. 1). Therefore, only bacteria that have lost the plasmid will grow. These observations indicated the feasibility of using inducible *mazF* expression as a method for counter-selection during double-crossover homologous recombination as depicted in Figure 2F. We successfully validated this approach by deleting the adhesin *fadA* in *F. nucleatum*, as evident from the absence of *fadA* mRNA on a northern blot (Fig. 2G). In combination, these five plasmid-based systems developed here provide much-needed genetic tools to accelerate functional genomics in *F. nucleatum* and likely other members of the understudied phylum of Fusobacteriota.

### The σ^E^ regulon in F. nucleatum

To define a potential σ^E^ stress response in *F. nucleatum*, we cloned the candidate fusobacterial *rpoE* gene C4N14_09830 into the inducible pVoPo-03 plasmid. We then used RNA-seq to determine the initial transcriptional response upon induction of this gene for 30 min during mid-exponential growth. Expression of C4N14_09830 did not affect bacterial growth at this time point (*SI Appendix*, Fig. S2); in fact, growth inhibition was observed only 2.5 h after induction. This indicates that aberrant activation of C4N14_09830 negatively affects cell growth only upon prolonged expression. Applying a false discovery rate (FDR) of ≤ 0.05, our global gene expression analysis identified 147 upregulated (log_2_ fold change ≥ 1) and 23 downregulated (log_2_ fold change ≤ -1) genes, as compared to empty vector control (Fig. 3A). The downregulated transcripts mostly encode membrane proteins and include an orthologue of the IM galactose transporter MglB and three similar multicistronic operons encoding envelope proteins, such as a FadA-domain containing protein, OmpA family proteins and type 5a autotransporters (Dataset S1).

**Figure 3:**
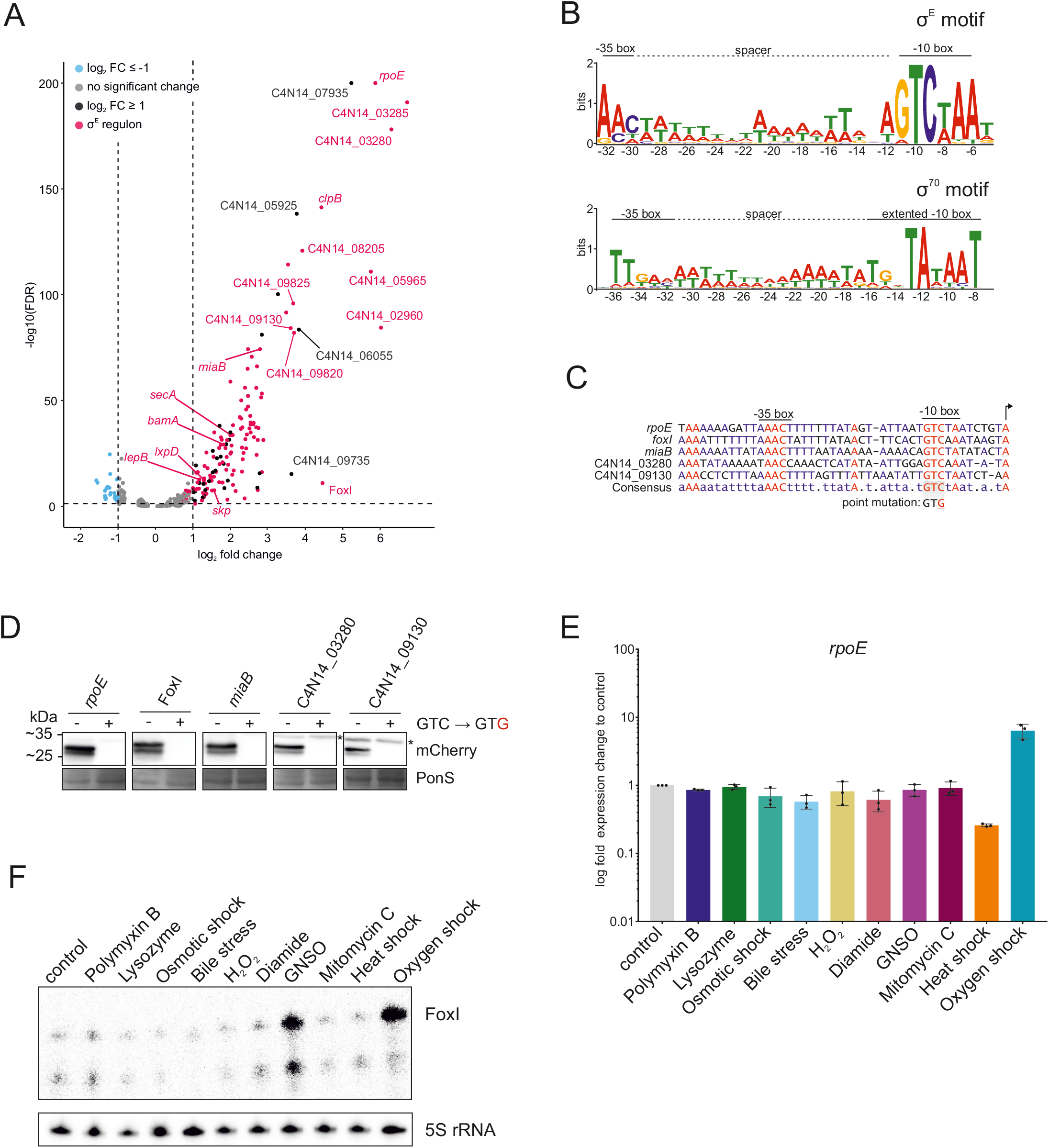
The σ^E^ regulon in *F. nucleatum*. (A) Volcano plot of the global gene expression changes in *F. nucleatum* after 30 min of σ^E^ induction. Gene expression of bacteria carrying pVoPo-03-σ^E^ is compared to cells carrying pVoPo-03 serving as empty vector control. Genes were considered significantly upregulated with a log_2_ fold change ≥ 1 (black) and significantly downregulated with a log_2_ fold change ≤ -1 (blue) with a false-discovery rate (FDR) ≤ 0.05 (dashed horizontal line). The σ^E^ regulon (red) includes all transcriptional units that harbor the identified σ^E^ binding motif in their promoter region. (B) Top, motif analysis via MEME (106) for all genes significantly upregulated upon σ^E^ induction. Transcriptional start sites (TSS) of all upregulated genes were manually annotated and 50 nt upstream of the identified TSSs were used as input for MEME. The conserved -10 and -35 boxes are indicated, as well as the AT-rich spacer in between both boxes. Bottom, the previously identified promoter motif for σ^70^ with an extended -10 box and a less pronounced -35 box (17). (C) Alignment of the promoter regions for selected genes identified as part of the σ^E^ regulon. A point mutation inserted into transcriptional reporter constructs (see (D)) is indicated. (D) Western blot analysis for mCherry expressed from transcriptional reporter plasmids harboring the native promoters (GTC) or a point mutation in the conserved -10 box (GTG) for selected genes shown in (C). Total proteins samples were collected during mid-exponential phase for western blot analysis. Ponceau S (PonS) staining is shown as loading control. Representative images of three independent experiments are shown. Unspecific bands are marked by an asterisk. (E) RT-qPCR analysis for *rpoE* mRNA after exposing *F. nucleatum* to the indicated stress conditions for 60 min. Data are normalized to the control and plotted as the average of three biological replicates with the standard deviation. (F) Northern blot probed for the sRNA FoxI using total RNA samples extracted from *F. nucleatum* treated with the indicated stress conditions for 60 min. The smaller band represents a degradation or degradation event. GNSO, *S*-nitrosoglutathione.

Analysis of the promoter regions of the upregulated genes revealed a common motif with a ‘GTCWAA’ in the -10 box and a less distinct ‘AAC’ in the -35 box separated by an AT-rich spacer region (Fig. 3B). This motif closely resembles the well-established consensus σ^E^ binding sites in *E. coli* or *P. aeruginosa* (61, 62). Additionally, the 18 nt-spacing between the transcriptional start site and the -10 box is very similar to *E. coli* (*SI Appendix*, Fig. S3) (31). Importantly, this motif is distinct from the previously identified σ^70^ binding site, in both the -10 and -35 regions (Fig. 3B). The putative σ^E^ motif is present in 28 transcriptional units consisting of 127 genes and accounted for 113 of the 144 upregulated genes (Dataset S1). Interestingly, we observed that in 14 cases σ^E^ activation initiated transcription of sub-operons (Dataset S1), leading to an uncoupling of gene expression from the upstream genes of these operons.

Besides the candidate *rpoE* gene itself, the two downstream genes in its operon (Fig. 1B) were also highly upregulated upon induction of this putative ECF (Fig. 3A), reflecting the established self-amplification of the σ^E^ response in *E. coli*, where σ^E^ directly activates its own promoter (63).

Despite the evolutionary distance of Fusobacteriota to Proteobacteria (Fig. 1A) (14), the transcriptional response described here exhibits several similar features to the σ^E^ regulon in *E. coli*. This includes, for example, upregulation of homologous genes important for the insertion of OMPs (*bamA*, *skp*) or lipid A biosynthesis (*lxpD*) (31, 33, 64–66). The observed target gene conservation classifies the candidate ECF protein C4N14_09830 as a σ^E^ homologue and thus we will refer to it as σ^E^ from here on. Interestingly, three genes (*ftsY*, *secA*, *lepB*) essential for SEC-dependent protein translocation across the IM (67, 68) were induced as well; none of them had previously been linked to σ^E^. 24 genes in the transcriptional response lack any functional prediction, including the most highly upregulated dicistronic operon C4N14_03280-C4N14_03285 (Fig. 3A). This raises the question if these genes are involved in envelope maintenance or protein translocation as well or if they represent an entire new function in the σ^E^-mediated stress response.

Recent studies have identified σ^E^-activated sRNAs in several different species (39-42, 49, 69, 70). Here, we observed a clear increase of the levels the oxygen-induced 87-nt sRNA FoxI upon induced expression of σ^E^ in *F. nucleatum* (17). This regulation suggests that the fusobacterial σ^E^ response might possess a noncoding arm, to which we will return below.

### Validation of σ^E^ target genes using transcriptional reporters

To confirm a subset of the identified σ^E^ target genes with an orthogonal method, we constructed five transcriptional reporters, in which the promoter regions of these targets, including the sRNA FoxI, drive mCherry expression. To confirm that the fusobacterial σ^E^ protein controls its own transcription, we included the promoter of the *rpoE* gene. All these promoters harbor in their -10 region a conserved cytosine shown to be critical for recognition by σ^E^ in *E. coli* (71) (Fig. 3C). Strikingly, a C-to-G point mutation at this position completely abolished transcription from these five selected fusobacterial promoters (Fig. 3D). These data indicate that these genes depend on σ^E^ for their transcriptional activation and support the relevance of the identified promoter motif for recognition by σ^E^.

### Oxygen-dependent activation of the σ^E^ response in F. nucleatum

It is well established that σ^E^ is activated in different bacterial species by various distinct stressors, such as unfolded proteins, osmotic stress, heat shock, singlet oxygen or oxidative stress (20, 29, 72, 73). To better understand what activates σ^E^ in *F. nucleatum*, we monitored *rpoE* mRNA levels upon exposure to different sources of envelope (polymyxin B; lysozyme; bile), osmotic (NaCl), and oxidative stress (H_2_O_2_; diamide; S-nitrosoglutathione (GNSO)); DNA damage (mitomycin C), heat shock (42°C) and oxygen exposure (Fig. 3E). Surprisingly, in this anaerobe bacterium, we observed a selective σ^E^ induction upon oxygen exposure. Importantly, the envelope-penetrating antibiotic polymyxin B, which is a well-established activator of σ^E^ in *E. coli*, did not induce *rpoE* in *F. nucleatum*, despite the fact that polymyxin B is active against this Gram-negative species (74).

FoxI, originally reported as an oxygen-responsive sRNA, is now found to possess a promoter that is stringently controlled by σ^E^ (Fig. 3C-D). Profiling FoxI expression in response to the full panel of stressors described above, we observed an almost selective increase in FoxI levels upon oxygen exposure (Fig. 3F, Supplementary Fig. 4), with the exception of nitrosative oxidative stress (GNSO) that induced this sRNA as well. Interestingly, *rpoE* mRNA levels did not increase after treatment with GNSO, suggesting that σ^E^ might not be the only regulator of FoxI (Fig. 3E-F). Supporting the strong induction of σ^E^ upon oxygen exposure, the transcripts of 4 additional genes of the σ^E^ regulon showed a similar increase after oxygen exposure in comparison to the untreated control (*SI Appendix*, Fig. S5).

*F. nucleatum* subspecies *nucleatum*, used in this study, harbors no additional ECFs. Yet, it does encode three conserved σ factors (*SI Appendix*, Fig. S6A): houskeeping RpoD; a second uncategeorized σ^70^-family member C4N14_05515; and a putative homolog of SigH of the Clostridiales. The strain we used also harbors a rare putative σ factor (C4N14_03400) found only in some members of Fusobacterales, Staphylococci, Clostridiaceae and on plasmids of Enterococcaceae. The alternative sigma factor RpoN is absent in *F. nucleatum* subspecies *nucleatum* strain used here. Testing a possible activation of the three conserved σ factors under different stress conditions by using RT-qPCR, we observed only mild expression changes upon oxygen exposure (*SI Appendix,* Fig. S6B). According to our RNA-seq data, these σ factor genes do not respond to σ^E^ either (Dataset S1), suggesting that σ^E^ is a main factor in the response to molecular oxygen.

### Global analysis of the oxygen response in F. nucleatum

In light of the specific activation of σ^E^ and FoxI by oxygen (Fig. 3E-F), we investigated the global activation of the σ^E^ regulon by exposing *F. nucleatum* to oxygen for 20 min, followed by RNA-seq analysis. We observed a total of 289 significantly regulated genes (FDR ≤ 0.05) with 174 upregulated (log_2_ fold change ≧ 1) and 115 downregulated genes (log_2_ fold change ≤ -1) (Fig. 4A). The upreguated genes included the *rpoE* operon and the sRNA FoxI confirming their activation by oxygen. 19 additional genes of the σ^E^ regulon identified in Fig. 3A were also upregulated (Fig. 4B), including the dicistronic operon C4N14_03280-C4N14_03285. Of note, sensing of oxygen by *F. nucleatum* induced differential expression of 15 transcription factors (Dataset S2), potentially indicating a wide-spread response beyond σ^E^.

**Figure 4:**
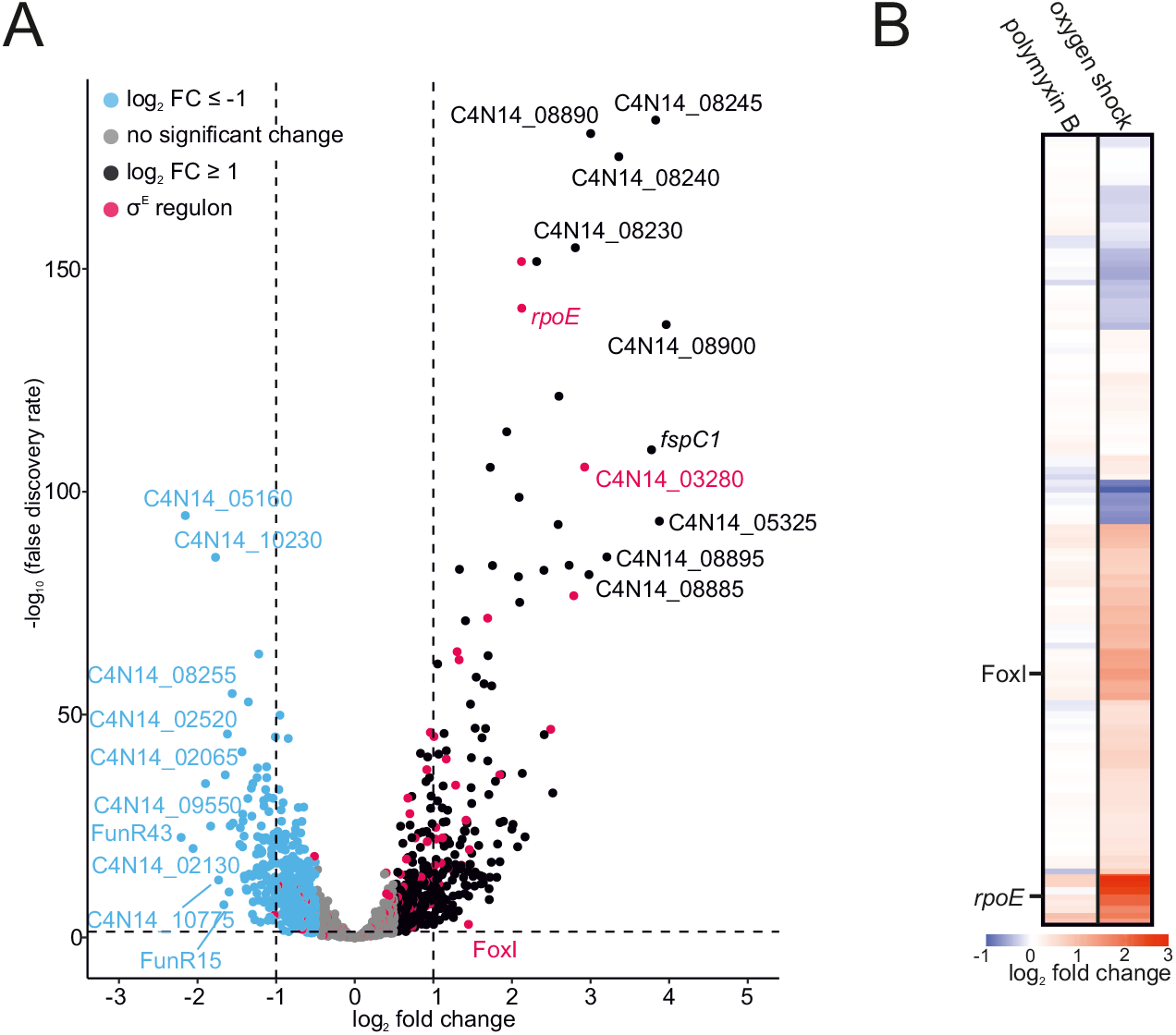
Transcriptional response of the anaerobe *F. nucleatum* to oxygen. (A) Volcano plot of the global gene expression changes in *F. nucleatum* after 20 min of oxygen exposure. Differential expression analysis was carried out by comparing the treated samples to an untreated control kept in the anaerobic chamber. Genes were considered significantly upregulated with a log_2_ fold change ≥ 1 (red) and significantly downregulated with a log_2_ fold change ≤ -1 (blue) with a false-discovery rate (FDR) ≤ 0.05 (dashed horizontal line). The σ^E^ regulon (red) contains all transcriptional units that harbor the identified binding motif in their promoter region. (B) Overview the gene expression changes for all members of the σ^E^ regulon upon exposure to 400 ng ml^-1^ polymyxin B or oxygen exposure for 20 min. The heatmap displays the log_2_ fold changes. The sRNA FoxI and rpoE are indicated.

### The sRNA FoxI is a negative regulator of the σ^E^ response

In previous work (17), we showed that the FoxI sRNA acted as a negative regulator of the abundant OM porin FomA. Although *fomA* did not pass our cutoff for significantly regulated transcripts (Fig. 3A), manual inspection of the global RNA-seq data revealed a clear decrease of *fomA* mRNA levels upon induced σ^E^ expression (log_2_ fold change of - 0.73). To test if FoxI might act as the negative regulator of σ^E^ in *F. nucleatum*, we used our *mazF-*based gene deletion tool pVoPo-04 to generate a Δ*foxI* strain (Fig. 5A). Following complementation of this strain with the inducible σ^E^ expression plasmid, we performed a global RNA-seq analysis after σ^E^ induction for 30 min. A comparison of downregulated genes in the wild-type (WT) versus and the Δ*foxI* strain showed that *fomA* mRNA levels were not reduced when FoxI was absent (Dataset S1). Similarly, the Δ*foxI* strain failed to downregulate *mglB* after σ^E^ expression (Fig. 5B), suggesting that the *mglB* mRNA might be another FoxI target. Nonetheless, *mglB* was the only other mRNA to show altered levels in the Δ*foxI* background, indicating that other targets of FoxI might be primarily regulated on the level of translation. Furthermore, additional σ^E^-dependent sRNAs that compensate for FoxI function might be present, similar to what is seen in *E. coli* (Gogol et al., 2011) and *V. cholerae* (Peschek et al., 2019).

**Figure 5:**
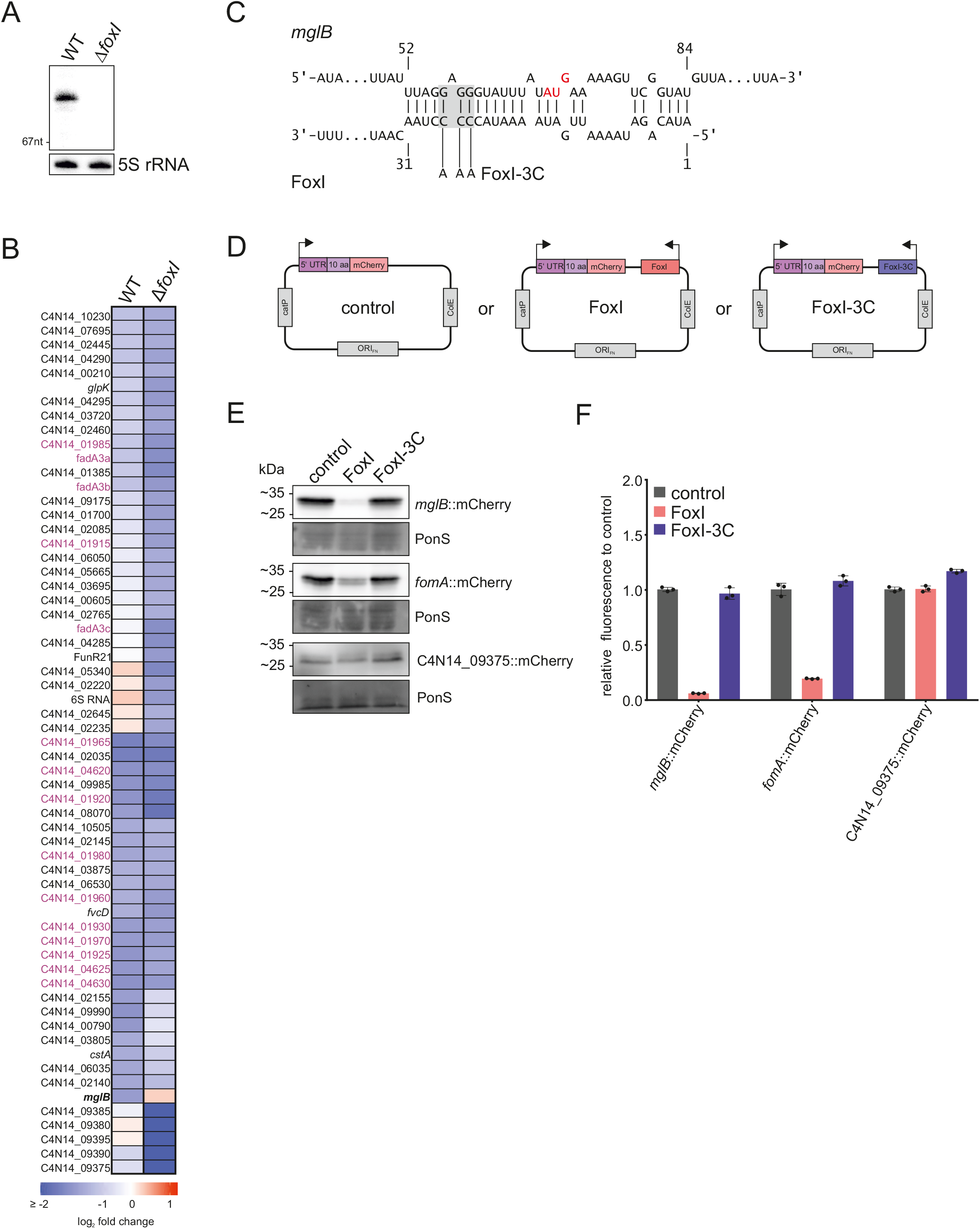
The sRNA FoxI as a negative regulator of the σ^E^ response. (A) Northern blot detection of FoxI using total RNA samples extracted from *F. nucleatum* wild-type (WT) or Δ*foxI* generated via the deletion system pVoPo-04. The 5S rRNA served as loading control. (B) Differential gene expression upon σ^E^ induction in wild-type (WT) *F. nucleatum* or in the FoxI deletion strain (Δ*foxI*). The heatmap displays log_2_ fold changes of genes that are significantly downregulated in either background (log_2_ fold change ≤ - 1; FDR ≤ 0.05). *mglB* is marked in bold as the only gene that is not downregulated in the Δ*foxI* background upon σ^E^ induction. Members of the three multicistronic operons started by FadA-domain containing genes are marked in purple. (C) Schematic representation of IntaRNA (93) prediction of base-pairing between *mglB* mRNA and FoxI. The AUG start codon of *mglB* is marked in red. The mutation of the sRNA for FoxI-3C is indicated in grey. (D) Schematic representation of the translational reporter constructs used in (E). *F. nucleatum* was either transformed with pVoPo-02 plasmids carrying mCherry fused to the 5’-region of the target gene only (control), or in combination with the expression cassette for FoxI (FoxI) or the seed region mutant FoxI-3C (FoxI-3C). (E) Representative western blots for each gene tested in the translational reporter system. C4N14_09375 served as control gene as it does not harbor any predicted FoxI-binding site. Ponceau S (PonS) staining is shown as loading control. (F) Quantification of mCherry by flow cytometry for the same constructs as shown in (E). The average of three biological replicates relative to that of the control (ctrl) is displayed together with the standard deviation.

### Post-transcriptional repression of envelope protein MglB by FoxI

An *in silico* prediction of RNA interactions indicated a potential binding site of the FoxI sRNA across the Shine-Dalgarno sequence of *mglB* (Fig. 5C), expected to prevent MglB synthesis upon binding of the sRNA. To validate the *mglB* mRNA as a FoxI target, we constructed a translational reporter based on plasmid pVoPo-02, expressing the 5’ UTR and the first 10 aa of *mglB* as a fusion to mCherry from a constitutive promoter. Subsequently, we added expression cassettes for FoxI or a Fox I mutant that carries mutations in its seed region (FoxI-3C) (Fig. 5D). Western blot analysis showed a strong reduction of the MglB::mCherry protein in the presence of FoxI compared to the control (Fig. 5E and *SI Appendix*, Fig. S7A), similarly to a mCherry fusion of the known target *fomA*. We further confirmed this effect by quantification of the fluorescence signal of the mCherry fusions via flow cytometry (Fig. 5F). These interactions are likely mediated via direct base-pairing since co-expression of the seed region mutant FoxI-3C did not downregulate these translational reporters (Fig. 5E-F). No regulation was observed with a C4N14_09375::mCherry fusion, chosen as a negative control since the C4N14_09375 mRNA does not harbor a predicted FoxI binding site. Combined, the results reveal *mglB* as a second target of the sRNA FoxI and highlight the application of our translational fusion system to validate sRNA-mediated mRNA regulation in *F. nucleatum*.

### Global transcriptome changes induced by FoxI expression and sRNA target identification

Induced overexpression of sRNAs is a powerful approach to capture the targetome of sRNAs (42, 75, 76), which includes successful experimental target searches for σ^E^-dependent sRNAs in Proteobacteria (37, 40, 42). Here, we took a similar approach and pulse-expressed FoxI, FoxI-3C as well as FoxI-C4A, a mutant that carries only a single point mutation in the seed region, in the *F. nucleatum* Δ*foxI* strain for 20 min. Of note, under these experimental conditions, FoxI did not affect bacterial growth (*SI Appendix*, Fig. S2). RNA-seq identified 30 downregulated mRNAs as potential FoxI targets (−0.5 ≤ log_2_ fold change ≥ 0.5; FDR ≤ 0.5); for most of these, repression was lost when expressing the seed region mutants FoxI-3C and FoxI-C4A (Fig. 6A, Dataset S3). The strongest negative regulation was observed for the C4N14_09375-C4N14_09395 operon of unknown function (Fig. 6A). However, this operon was also downregulated by expression of the FoxI seed mutants, arguing against it being a direct target of FoxI. More importantly, we observed downregulation of the FoxI target *mglB* and of *mglA*, the gene immediately downstream of *mglB* in the *mglBAC* operon. Curiously, pulse-expression of FoxI-C4A also reduced the *mglB* transcript levels but did not affect *fomA* mRNA (Fig. 6A), possibly indicating a more robust target interaction of FoxI with *mglB* in comparison to *fomA* (Supplementary Fig. 8). In addition, several genes of the σ^E^ regulon, including the σ^E^ operon itself, were downregulated upon expression of a WT copy of FoxI, but not the seed mutants (Fig. 6A; purple). Interestingly, *in silico* target predictions for FoxI only revealed poor binding sites for these genes (Dataset S4), suggesting that some of them are regulated as an indirect consequence of FoxI-mediated relief of basal activation of σ^E^. Overall, our analysis supports the identification of *mglB* as a direct target of the sRNA. Nevertheless, the moderate changes in RNA levels upon FoxI expression suggests that this sRNA might act primarily on the translational level, similar to the sRNA Spot 42 in *E. coli* which blocks translation within the *galETKM* mRNA without affecting mRNA levels (77).

**Figure 6:**
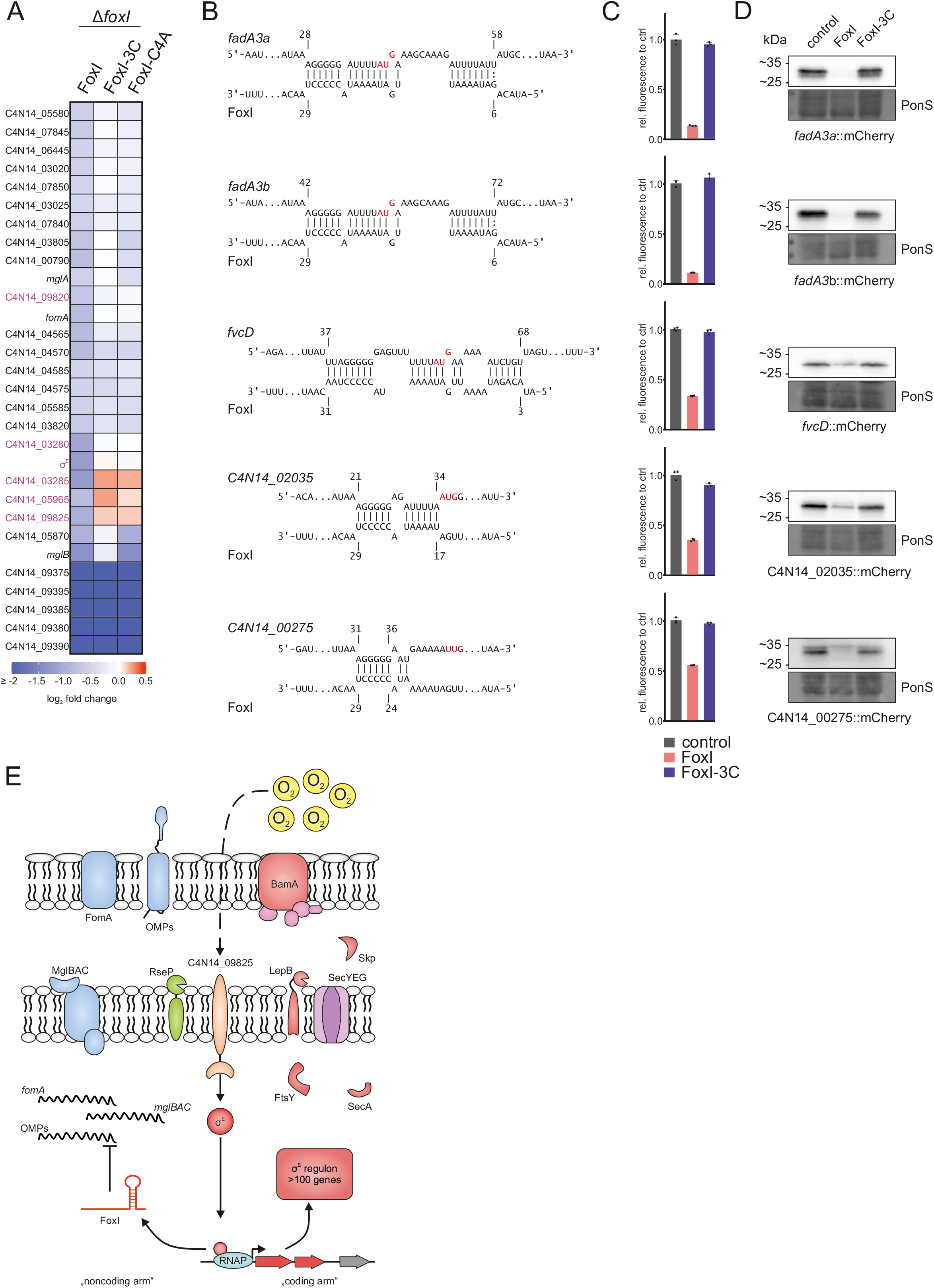
The full target spectrum of the sRNA FoxI. (A) Overview of all significantly downregulated genes (log_2_ fold change ≤ -0.5; FDR ≤ 0.05) upon pulse expression of FoxI or the seed region mutants FoxI-3C or FoxI-C4A in the Δ*foxI* background. The heatmap displays the log_2_ fold changes. Genes of the identified σ^E^ regulon (see Figure 3) are marked in purple. (B) Schematic representation of IntaRNA target predictions between different target mRNAs and FoxI. The AUG start codons are marked in red. (C) Quantification of mCherry by flow cytometry for the translational reporter plasmids carrying the target 5’ region shown in (B) alone (control) or in the presence of FoxI or FoxI-3C. The average of three biological replicates relative to that of the control is displayed together with the standard deviation. (D) Representative images of western blots of total protein samples for bacteria expressing the indicated reported constructs probed for mCherry. Ponceau S (PonS) staining is shown as loading control. (E) Model of the σ^E^ regulon in *F. nucleatum*. σ^E^ is released from its putative anti-sigma factor (C4N14_09825) and upregulates expression of its regulon consisting of > 100 genes (marked in red). This includes *bamA* and *skp*, important for insertion of OMPs as well as *lepB*, *ftsY* or *secA*, which are involved in protein-translocation across the inner membrane. σ^E^ also activates the transcription of the sRNA FoxI. FoxI, in turn, downregulates membrane-associated proteins such as the IMP complex *mglBAC* and the OMP *fomA* as well as other OMPs.

### Evidence for multi-target regulation by the FoxI sRNA

In *E. coli*, the σ^E^-dependent sRNAs MicA, RybB and MicL repress the mRNAs of several OMPs as well as the abundant Lpp protein (37, 38, 42). Since unfolded OMPs and Lpp are potent triggers of σ^E^, the sRNA-mediated translational inhibition results in a negative feedback-loop within the σ^E^ response. Interestingly, we observed repression for several OMPs upon σ^E^ induction in *F. nucleatum* (Fig. 3) and *in silico* target prediction indicates promising binding sites for FoxI for some of these OMPs (Fig. 6B; Dataset S3). We therefore hypothesized that FoxI might block the synthesis of multiple OMPs without directly affecting their RNA levels. To test this, we applied our translational reporter system to five additional candidate target mRNAs, whose levels decreased upon σ^E^ induction: *fadA*2, *fadA*3, C4N14_00275, C4N14_02035, and C4N14_08545. In all these cases we observed a strong translational repression upon constitutive FoxI expression, but not upon expression of the FoxI-3C mutant measured by flow cytometry (Fig. 6C) and western blot analysis (Fig. 6D, *SI Appendix* Fig. S7B). Thus, we have obtained evidence for FoxI-mediated post-transcriptional repression of at least seven mRNAs of envelope proteins, lending further support to a model in which the conserved FoxI sRNA acts as the global non-coding arm of the fusobacterial σ^E^ response (Fig. 6E).

## DISCUSSION

Despite the growing appreciation of their importance for health and disease, the vast majority of the >4,500 bacterial species that constitute the human microbiota are currently molecular *terra incognita* (78). Research in this area is hampered by the fact that genetic modification of these bacteria is notoriously difficult and thus we lack the ability to genetically dissect their physiology and molecular principles of gene regulation. *F. nucleatum* has emerged as a new paradigm for such microbes and its ability to colonize distal body sites has increasingly been recognized as a medical problem (2). Here, we present a broad suite of genetic tools for *F. nucleatum* and apply these to uncover a conserved stress response mediated by the ECF σ^E^ in this bacterium, which was triggered by oxygen instead of general envelope perturbation.

Our present knowledge of the architecture of the σ^E^ response primarily stems from studies in a few γ-proteobacterial species (30), were it was shown to be a central player in combatting envelope stress. σ^E^ upregulates factors that ensure proper folding and insertion of OMPs or lipoproteins and thereby helps to maintain and shape the bacterial envelope. Since *F. nucleatum* exhibits a very large phylogenetic distance to these proteobacterial species (14), it was surprising to discover in *F. nucleatum* a σ^E^ regulon of similar architecture to *E. coli* (31, 38): many of the genes controlled by σ^E^ possess envelope-related functions, and there is a noncoding arm, i.e. the FoxI sRNA, many of whose target mRNAs also encode proteins with envelope-related functions. Strikingly, however, σ^E^ itself is not induced by known triggers of envelope stress but by exposure to oxygen. While the molecular mechanism of oxygen-mediated activation remains to be elucidated, it is tempting to speculate that σ^E^ in this early-branching anaerobic bacterium serves the role of an environmental sensor, while sharing the envelope remodeling function with *E. coli* σ^E^.

As to specific σ^E^-controlled genes, *skp* (79) and *bamA* (66) encode proteins that work cooperatively to ensure proper insertion of unfolded OMPs into the outer membrane. The protein encoded by *lpxD*, on the other hand, is integral for LPS biosynthesis (80) – another important component of the envelope of Gram-negative bacteria. We also identified a consensus σ^E^-binding site, consisting of the conserved -10 and -35 boxes, similar to *E. coli* and *P. aeruginosa* (31, 62). In light of the evolutionary distance between Fusobacteriota and other bacterial phyla (14), this conservation suggests that the σ^E^ response represents a very deeply rooted regulon that maintains bacterial envelope homeostasis. The σ^E^ regulon in *F. nucleatum* also includes genes encoding three integral members of the SEC-dependent protein translocation pathway: *ftsY*, *secA* and *lepB*. The FtsY and SecA proteins facilitate the translocation process, in a co-translational or post-translational manner, respectively. The signal peptidase LepB releases translocated proteins into the periplasm (81). Interestingly, these proteins have not been found to be under σ^E^ control in other bacteria, but increased translocation capacity would be expected to act in synergy with enhanced OMP insertion. After all, the SEC-pathway is the major transport mechanisms for OMPs to the periplasm (81, 82). Although we still lack a functional understanding of the 117 genes that are part of the σ^E^ regulon in *F. nucleatum*, it is clear that at least part of the physiological role of σ^E^ in this anaerobic bacterium is envelope maintenance, reflecting a core function for this ECF.

The link between σ^E^ and the response to molecular oxygen is supported by the common activation of 23 genes upon σ^E^ induction and oxygen exposure (Fig. 4A-B). An interesting example is activation of the dicistronic operon *ccdA*-*msrAB*, which is paralogues to an operon previously linked to the defense against hydrogen peroxide in *F. nucleatum* (Dataset S1) (13). Thus, the σ^E^ regulon in *F. nucleatum* might serve a dual function by neutralizing oxygen and by modulating the bacterial envelope, acting in synergy in this anaerobe bacterium. However, based on the differential expression of 15 transcription factors it is clear that σ^E^ is not the only mediator of an oxygen-induced response. Therefore, it will be important to understand how this anaerobe senses oxygen and transmits the signal to activate σ^E^. As to the actual activation mechanism, we note that *F. nucleatum* has no homologue of the protease DegS, which senses unfolded OMPs in *E. coli* (83). DegS initiates a proteolytic cascade that leads to the degradation of the anti-sigma factor and release of σ^E^ (84, 85). The lack of DegS might imply that *F. nucleatum* uses alternative ways of perceiving and relaying stress signals that lead to σ^E^ activation. These might involve phosphorylation as shown for the *Vibrio parahaemolyticus* σ factor EcfP (27) or a TCS as seen during activation of SigE in *Streptomyces coelicolor* (28). Interestingly, the fusobacterial TCS ModRS was recently shown to be involved in the response to H_2_O_2_ (13). However, ModRS is not activated by oxygen (Dataset S3). Alternatively, the anti-sigma could be directly involved in sensing oxygen and subsequently trigger σ^E^ activation. Such a mechanism has been shown for singlet oxygen in *Rhodobacter sphaeroide*s, where the anti-sigma factor ChrR directly responds to the reactive oxygen species and releases σ^E^ (21, 24, 86). Another possibility is activation via co-factors, such as [4Fe-S]^2+^ or heme, which present widespread oxygen-sensing mechanisms in bacteria (87).

One of the most striking findings of the σ^E^ response of *F. nucleatum* is the conservation of a noncoding repressor arm, constituted by the sRNA FoxI, in an organism that lacks known sRNA chaperones. Although sRNAs are frequently part of regulatory circuits, there is only one example of broad conservation in different phyla: the sRNAs controlled by the iron uptake regulator Fur (88). Fur is present in Gram-positive and Gram-negative bacteria because iron is essential for all bacteria (89). The Fur-dependent sRNAs, such as RyhB in *E. coli* (90), expand Fur’s transcriptional repressor function upon iron starvation (91, 92). Similarly, sRNAs form the repressive arm of the σ^E^ response and play an important role in downregulating envelope proteins (38). However, evidence for a conservation of this function has come from the single phylum of Proteobacteria, i.e., from *E. coli* and *Salmonella* (39, 42, 49, 69), *Pseudomonas aeruginosa* (70) and *Vibrio cholerae* (40, 41). Here, we find that the sRNA FoxI is expressed in a σ^E^-dependent fashion and represses the translation of the OM porin FomA, the MglBAC galactose-uptake system and leading genes of operons that encode type 5a autotransporters, another class of abundant fusobacterial OMPs (12, 93–95). Our observations suggest that FoxI reduces translation of membrane proteins, thereby limiting the burden on the Sec- and BAM-dependent membrane protein insertion pathways. This is likely also reflected in the observation that expression of FoxI in the Δ*foxI* background decreases expression of the *rpoE* and C4N14_03280-C4N14-03285 operons (Fig. 6A), which might indicate a decreased activation of the basal σ^E^ stress response. FoxI and σ^E^ thus work synergistically to maintain envelope homeostasis and integrity in *F. nucleatum* (Fig. 6E), mirroring the coding arm and noncoding arm principle established by σ^E^-dependent sRNAs in *E. coli* (38). Nevertheless, the fact that we did not observe a strict dependence of σ^E^-repressed targets on FoxI suggests that there could be additional σ^E^-dependent sRNA. This has been seen in *E. coli* (MicA, RybB) (38) or *V. cholerae* (VrrA, MicV) (40), where two sRNAs share common targets and compensate for another.

As mentioned, all σ^E^-dependent sRNAs studied thus far rely on the RNA chaperone Hfq for their activity (42, 46, 49), but *F. nucleatum* lacks known sRNA chaperones. It nevertheless remains possible that a yet unidentified RNA-binding protein (RBP) plays a role in sRNA-mediated regulation in *F. nucleatum*. To facilitate the discovery of such RBPs, proteins interacting with FoxI could be identified using the sRNA as bait for pulldown experiments (76). A promising candidate for this role might be the RBP KhpB, which has been found to bind RNA in *Streptococcus pneumonia* and *Clostridium difficile* (96–98), and for which *F. nucleatum* harbors a homologue.

Overall, our study highlights the conservation of the regulatory principle of the bacterial σ^E^ response, despite the evolutionary distance of Fusobacteria to other bacterial clades and provides much-needed tools to dissect *F. nucleatum* gene function in order to accelerate research into this clinically-relevant bacterium.

## MATERIAL AND METHODS

### Strains and growth conditions

All oligonucleotides, plasmids or strains used in the present study can be found in Dataset S5. *Fusobacterium nucleatum* subspecies *nucleatum* ATCC 23726 (*F. nucleatum*) was acquired from the American Type Culture Collection (ATCC). *F. nucleatum* was routinely grown at 37 °C in 80:10:10 (N2:H2:CO2) on 2% agar BHI-C plates (brain–heart infusion (BHI), 1% (w:v) yeast extract, 1% (w:v) glucose, 5 µg ml−1 of hemin; 1% (v:v) fetal bovine serum). For liquid growth Columbia broth medium was utilized. For selecting *F. nucleatum* carrying a plasmid or for selection steps during gene deletion, BHI-C agar plates were supplemented with 5 µg ml^-1^ thiamphenicol and liquid cultures with 2.5 µg ml^-1^ of the antibiotic. All solutions or plates were always pre-reduced overnight prior to use in the anaerobic chamber to ensure the absence of entrapped oxygen. For growing *F. nucleatum*, pre-cultures were prepared 24 h prior to inoculating the working cultures (1:50 dilution).

### Construction of improved backbone pVoPo-00

Our previously generated plasmid pEcoFus (17) was optimized and reduced in size as follows: The ColE replication of origin for *E. coli* as well as the *catP* resistance cassette for chloramphenicol/thiamphenicol were amplified from pEcoFus using primer pairs JVO-18069/ JVO-18070 and JVO-18071/ JVO-18072, respectively. The first primer pair includes a multiple cloning site and both fragments were assembled using NEBuilder Hifi Assembly Cloning kit (New England Biolabs) together with the promoter and 5’ UTR of the constitutive expressed flavodoxin C4N14_09865 amplified from genomic DNA of *F. nucleatum* using JVO-18073/ JVO-18074. This yielded pFP76, which contains the *catP* gene constitutively, expressed from the inserted fusobacterial promoter. pFP76 was digested with PvuI and NotI and ligated with a similarly digested origin of replication for *F. nucleatum* (F-ORI) excised from pEcoFus. This resulted in the improved *F. nucleatum* – *E. coli* shuttle vector pVoPo-00.

### Construction of transcriptional reporter system pVoPo-01

The here generated pVoPo-00 was opened via inverted PCR (JVO-18273/ JVO-18274) and used for an assembly reaction together with mCherry amplified from pDSW1728 (99) (JVO-18339/ JVO-18340) as well as a single strand oligonucleotide (ssOligo) JVO-18338 forming the 5’ UTR of *acpP* in *F. nucleatum*. The resulting vector was opened once more (JVO-18341/ JVO-18342) to insert a 50 bp promoter region of the constitutively expressed *accD* (C4N14_10115; ssOligo JVO-18344). This resulted in pVoPo-01, which was used to introduce the different promoter regions tested in this work. In all cases, pVoPo-03 was opened by inverted PCR (JVO-18341/ JVO-18342) and assembled with the different ssOligo containing the different promoter sites (Dataset S4).

### Construction of translation reporter system pVoPo-02

The vector pVoPo-01 was opened by inverted PCR (JVO-19090/ JVO-19091) and assembled with an ssOligo (JVO-20214) to include a ScaI restriction site at transcriptional start and an in-frame XhoI site with the coding sequence of mCherry. This yielded pVoPo-02, which was used to generate translational fusions (see below).

### Construction of inducible system pVoPo-03

To construct an inducible expression plasmid, the *tetR*-GUS cassette from pRPF185 (100) was amplified (JVO-18371/JVO-18372) and assembled into the SpeI site of pVoPo-00. The resulting vector was opened via inverse PCR to remove the GUS gene and insert an XhoI restriction site (JVO-17537/ JVO-17538). The vector can be opened at transcriptional start site of the TetR-dependent promoter via the BamHI digestion.

### Construction of gene deletion system pVoPo-04

pVoPo-03 was digested with XhoI and BamHI. The *mazF* gene was amplified to include an XhoI and BamHI restriction site as well as its natural 5’ UTR. Both fragments were ligated. To generate the fusobacterial suicide vector pVoPo-04, the *tetR*-*mazF* expression cassette was transferred into pFP76 via the SpeI site (JVO-18075/ JVO-18076).

### Construction of translational fusions for studying the post-transcriptional regulation mediated by FoxI

pVoPo-02 was further digested with EcoRI to insert the FoxI or FoxI-3C overexpression cassette from pFP10 and pFP70, respectively, via isothermal assembly reaction to ensure the desire directionality. Each vector was then digested with ScaI and XhoI and ligated with similarly digested PCR products and regions of interest containing the 5’ UTR and the first 30 nucleotides of the target genes, generating the target vectors (see Dataset S4 for primers used).

### Electroporation of F. nucleatum

Preparation of electro-competent cells as well as delivery of plasmid DNA was achieved as described previously (17). In short, cells were harvested from the mid-exponential phase, pelleted and washed five times with ice-cold pre-reduced 10 % (v:v) glycerol solution. 5 OD cells per transformation were used to transform 100 ng (replicative plasmid) or 10 µg (suicide vectors; dialyzed) DNA via electroporation (2.0 kV, 1-mm gap). Bacteria were recovered for 2 h prior to plating on BHI-C plates containing 5 µg ml^-1^ of thiamphenicol enabling selection of successful transformants.

### Evaluation of mazF expression for counter-selection

To create a *mazF*-containing plasmid that allows replication in *F. nucleatum*, the F-ORI was inserted into the PvuI and NotI site of pVoPo-02 generating pVoPo-01-*mazF* (p.*mazF*). Subsequently, bacteria carrying p.*mazF* or the control vector pVoPo-01 (p.empty) used to prepare pre-cultures in biological triplicates. For analysis of bacterial survival, the cells were grown to mid-exponential phase and exposed to either 100 ng ml^-1^ ATc or left untreated. Serial dilution were plated at 0 min, 30 min or 3 h after treatment. For this 10 µl were spotted in technical triplicates on BHI-C plates. Three days later, all technical replicates for the appropriate dilutions were counted and averaged for each biological replicate. The average and standard deviation is shown in Figure 2C. For analysis of growth, pre-cultures for three replicates were diluted as described above either in the presence of 100 ng ml^-1^ ATc or left untreated. Growth was monitored for 24 h using a plate reader and reported as the average of three biological replicates for each group in Figure 2D.

### Generating clean deletion mutant using pVoPo-04 system

To allow homologous recombination, 1 kB flanking up- and downstream of the target gene were amplified from genomic DNA of *F. nucleatum* and assembled in pVoPo-04 as described above. Transformation was carried out as described above and successful integration events were re-streaked on fresh BHI-C plates containing thiamphenicol. Colonies that grew had successfully integrated the suicide vector into the genome (marked as 1^st^ recombination). A single colony was used to inoculate an overnight culture in Columbia broth without selection pressure to allow the second recombination step to take place. The next day, the culture was diluted 1:50 into media containing 100 ng ml^-1^ anhydrotetracycline (ATc). This allows for the counter-selection due to the toxic expression of the toxin *mazF* and only bacteria having lost the plasmid can grow. After 4 h, serial dilutions were plated on BHI-C plates. The loss of the plasmid was verified via re-streaking resulting colonies on BHI-C and BHI-C containing thiamphenicol plates. Only colonies growing on BHI-C but not the plates with antibiotic were used for further validation via PCR to check for the loss of the target gene or reversion to the wild-type.

### Construction of protein and sRNA inducible expression vectors using pVoPo-03

To evaluate the effect of σ^E^ expression, we amplified *rpoE* from the genomic DNA of *F. nucleatum* and placed it in the BamHI site in pVoPo-03. To enable higher translation, we added a short synthetic 5’ UTR (101) by opening the vector via inverse PCR (JVO-19601/ JVO-18355) and assembling it with the ssOligo (JVO-19605). This generated pVoPo-03-σ^E^ (p.σ^E^). To achieve inducible expression of mCherry, we amplified the *mCherry* gene from pVoPo-01 (JVO-19865/JVO-19866) and replaced σ^E^ in p.σ^E^ through inverse PCR (JVO-18355/JVO-18346). To insert FoxI, FoxI-3C or FoxI-C4A under control of TetR, we amplified *foxI* from gDNA of *F. nucleatum*, FoxI-3C from pFP70 and FoxI-C4A from pFP181 and placed in the BamHI site of pVoPo-03 via isothermal assembly to have the sRNA start at the transcriptional site of the system.

### Construction of pVoPo-FP for the expression of different fluorescence proteins

Codon-optimized sequences of superfolder GFP (sfGFP), mNeonGreen and mScarlet-I were synthesized by Eurofins. The sequences can be found in Dataset S5. First, p.σ^E^ used as a template to insert mScarlet-I downstream of the 5’ UTR by opening the vector with JVO-19859 / JVO-19860. This product was used together with mScarlet-I amplified via JVO-19869 / JVO-19870 for an assembly reaction. The subsequent vector was opened with JVO-20852 / JVO-20853 and used with the ssOligo JVO-20859 in an assembly reaction to insert the promoter of the fusobacterial *acpP*. This resulted into pVoPo-mSc. To insert sfGFP and mCherry, the individual gene products were amplified with JVO-21080 / JVO-21081 and JVO-21082 / JVO-21083, respectively, and assembled with pVoPo-mSc opened via JVO-21079 / JVO-19859. This resulted in pVoPo-GFP and pVoPo-mCh. As the same strategy was unsuccessful for the generating a vector expressing mNeonGreen, we placed mNeonGreen in the backbone of pVoPo-02. For this, pVoPo-02 was opened with JVO-19275 / JVO-19276 and assembled with mNeonGreen amplified using JVO-19279 / JVO-19280. This yielded pVoPo-mNG. Of note, due to the difference in codon usage between *E. coli* and *F. nucleatum*, high expression of any gene, codon-optimized for or native to *F. nucleatum*, can result in spontaneous mutation or otherwise inactivation of the gene product in *E. coli*.

### Northern blot

Detection of RNA via northern blot was carried out as described before (17). In short, 3 µg of DNaseI treated total RNA was separated on a 6% polyacrylamide gel (7 M urea). Afterwards, the RNA was transferred to Hybond-XL membranes and hybridized overnight at 42 °C with [γ^32^]-ATP end-labelled deoxyribonucleotide probes (Dataset S4). The signal was visualized using a Typhoon FLA 7000 phosphoimager (GE Healthcare).

### Western blot

Detection of proteins using western blot was carried as described before (17). Briefly, 0.2 OD_600 nm_ units were loaded on denaturing SDS-polyacrylamide gel for SDS PAGE analysis. Proteins were transferred to polyvinylidene fluoride (PVDF) membranes. After transfer, equal loading was verified by staining with Ponceau S solution. mCherry was detected using rabbit anti-mCherry polyclonal antibody (Life Technology PA534974) as a primary antibody in combination with an anti-rabbit secondary antibody (Thermo Fisher Scientific, catalogue no. 31460).

### Sample collection for RNA-seq of σ^E^ expression

Three biological replicates for each group (wild-type: p.empty; p.σ^E^ and Δ*foxI*: p.empty; p.σ^E^) were grown to mid-exponential phase. All samples were induced with 100 ng ml^-1^ of ATc for 30 min. Samples were fixed by adding STOP Mix (95% (v:v) EtOH; 5% (v:v) phenol) and then snap-frozen in liquid nitrogen. Samples were stored at -80 °C until further processing. The Hot Phenol was used for RNA extraction as reported previously (17).

### Analysis of fluorescent protein expression by confocal microscopy

*F. nucleatum* carrying the individual pVoPo-FP plasmids were grown to mid-exponential phase. All following steps were conducted outside the anaerobic chamber. One ml of each culture was spun down and washed once in PBS. Next, the bacteria were fixed in 4% (w/V) PFA for 20 min at 4 °C. Afterwards, the cells were washed in once with PBS prior to overnight incubation in PBS at 4 °C. This step ensures proper maturation of the fluorescent proteins. The next day, the samples were imaged on ibdi_®_ chambered coverslips performed on a Leica SP5 laser scanning confocal microscope (Leica Microsystems) acquiring the fluorescence signal at the indicated wave lengths.

### Exposure of F. nucleatum to different stress conditions

Three biological replicates of *F. nucleatum* were grown to mid-exponential phase. The cultures were split into 3 ml aliquots and treated by adding a 1 ml of a 4x solution in Columbia broth of the following conditions for 60 min: polymyxin B (400 ng ml-1); lysozyme (125 µg ml-1); NaCl (600 mM); bile (0.05 % (w/V); H_2_O_2_ (400 µM); diamide (125 µM); S-nitrosoglutathion (GNSO, 250 µM); mitomycin C (625 ng ml-1). The final concentrations used are given. For the heat shock, the samples were placed in an incubator at 42 °C for the duration of the treatment. Regarding the oxygen exposure, the samples were poured into a petri dish and placed in an incubator at 37 °C outside of the anaerobic chamber for the duration of the treatment. One ml of Columbia broth was added as a control to untreated to the control samples. After 60 min samples were fixed through the addition of STOP mix and RNA extracted via the Hot Phenol protocol as mentioned above.

### Gene expression analysis via RT-qPCR

One µg of DNase-digested RNA was used as input to generate cDNA using the M-MLV reverse transcriptase (ThermoFisher Scientific) and random hexamer primers following the manufacturer’s instruction. The equivalent of 10 ng RNA was used for qPCR analysis using gene specific primers (Dataset S5). For this, the Takyon™ Master Mix was used according to the manufacturer’s protocol. The relative fold changes to the control were calculated based upon the 2^-ΔΔCt^ method (PMID: 11846609). The 5S rRNA was used as reference gene.

### Sample collection and analysis for translation fusion experiments by western blot

*F. nucleatum* carrying the individual translational fusion alone, or in combination with FoxI/ FoxI-3C was grown to mid-exponential phase and cells were quickly spun down and snap-frozen in liquid nitrogen. Thawed cells were resuspended in protein loading buffer and 0.2 OD_600 nm_ units were used for western blot analysis. Quantification of the fusion products signal in the western blot was carried out using Image J (102). Three biological replicates were analyzed in each case and the average together with the standard deviation reported.

### Sample collection and analysis for translation fusion experiments by flow cytometry

*F. nucleatum* carrying the individual translational fusion alone, or in combination with FoxI/ FoxI-3C was grown to mid-exponential phase and cells spun down for 3 min at 4.000 xg. This and all following steps were conducted outside of the anaerobic chamber. After removing the supernatant, the bacteria were fixated in 4% (w/V) PFA for 20 min at 4 °C. Next, the cells were washed with 1x PBS before incubating them with DAPI in PBS (100ng ml^-1^) for 5 min at room temperature. The bacteria were washed once more with PBS and resuspended in PBS. To ensure full maturation of the fluorescent protein the samples were left overnight at 4 °C. The next day, the fluorescence intensity was measured by flow cytometry at 615-620 nm for 50.000 cells of each sample determined by a DAPI^+^ signal.

### Sample collection and analysis for transcriptional reporter experiments

*F. nucleatum* carrying the individual transcriptional reporters was grown to mid-exponential phase as σ^E^ shows. At this time, samples were collected and snap-frozen. No quantification was carried out as no signal could be detected for samples with the point mutation.

### Sample collection for RNA-seq of sRNA expression

Three biological replicates for each group (Δf*oxI*: p.empty; FoxI; FoxI-3C; FoxI-C4A) were grown to mid-exponential phase. All samples were induced with 100 ng ml^-1^ of ATc for 20 min. Samples were treated as described above prior to performing RNA extraction according to the Hot Phenol protocol.

### Sample collection for RNA-seq upon oxygen and polymyxin B exposure

Three biological replicates of wild-type *F. nucleatum* were grown to mid-exponential growth phase. The cultures were exposed either to atmospheric oxygen concentrations outside of the anaerobic chamber (maintaining 37 °C) or treated with 400 ng ml^-1^ of polymyxin B for 20 min. Untreated samples were used as control. RNA extraction was performed according to the Hot Phenol protocol.

### cDNA library preparation for RNA-seq

The cDNA library preparation was carried out by Vertis (Munich, Germany). The first step consisted in rRNA removal. The rRNA depleted RNA was then fragmented via ultrasound (1 pulse of 30 seconds; 4 °C). The fragmented RNA was used to ligate an adapter to the 3’ end of the molecules. For first-strand cDNA synthesis, the M-MLV reverse transcriptase was used in conjunction with the introduced 3’-adapter serving as a primer. After purification, 5’ Illumina TruSeq sequencing adapters were ligated to the cDNA. The resulting cDNA was used as input for PCR amplification (10 – 20 ng µl-1). The amplified cDNA was purified using the Agencourt AMPure XP kit (Beckman Coulter Genomics) and evaluated via capillary electrophoresis. The cDNA was pooled and further purified to include only cDNA from 200 – 600 bp using a preparative agarose gel. The finished pooled libraries were sequenced by the Core unit SysMed (University of Würzburg) using an Illumina NextSeq 500 system and 75 bp read length.

### Read mapping and differential gene expression analysis

Reads from the RNA-seq experiments were trimmed and filtered using the FASTX toolkit (v.0.10.1; http://hannonlab.cshl.edu/fastx_toolkit). Mapping was performed using READemption (v.1.01) (103) against the genome sequence for *F. nucleatum* subsp. *nucleatum* ATCC 23726 (NZ_CP028109.1) downloaded from the National Center for Biotechnology Information (NCBI). Differential gene expression analysis was performed using DEseq2 (v.1.18.01) (104). The data from biological triplicates was used for the analysis. For the experiment concerning the σ^E^ expression, exposure to oxygen or polymyxin B, the genes had to show a log_2_ fold change ≤ -1 or ≥ 1 to be marked as significantly regulated. For the experiment concerning the pulse expression of FoxI or seed region mutants of the sRNA, the fold change of genes to be considered significantly changed was set to either ≤ -0.5 or ≥ 0.5. In both cases, the differentially expressed genes were only considered if their false-discovery-rate (FDR) was ≤ 0.05.

### Generation of phylogenetic tree

We obtained the data of the phylogenetic tree from *Coleman et al.* (2021) (14) and used it as input to visualize the phylogenetic tree via ggtree (105) to generate Figure 1A.

### Promoter analysis

For the identification of promoter motif, each significantly upregulated gene resulting from the σ^E^ expression were grouped into transcriptional units (see Fig. 3A). Based on our previous data for the transcriptional start sites (TSS) (17), we then extracted 50 nt upstream of each TSS for the individual transcriptional unit. Internal as well as secondary start sites of genes where we observed strong RNA-seq upregulation were treated the same. These nucleotide sequences were used as input for analysis via MEME (v.4.12.0) (106).

### In silico target prediction

Prediction of sRNA targets was carried using IntaRNA (v. 2.0.4) (107). As input we used all genes significantly downregulated with σ^E^ expression (see Fig. 3). For this, we extracted the nucleotide sequences for the entire CDS as well as the 5’ UTR. In case we were not able to determine the 5’ UTR, we took additional 50 nt upstream of the start codon. The prediction was carried out using standard settings with the exception of running it in the heuristic mode and allowing the seed region to consist out of up to three base pairs.

## Data and material availability

RNA-seq data can be accessed at NCBI’s GEO (https://www.ncbi.nlm.nih.gov/geo) under the accession no. GSE192339. Plasmids pVoPo-FP (with the individual fluorescence proteins) and pVoPo-01 to 04 have been deposited with Addgene (see Dataset S5).

## Author Contributions

F.P. performed most of the experiments. F.P., Y.Z. and V.C. constructed the expression vectors and performed the stress conditions experiments. F.P. performed data analysis. F.P. and J.V. designed research. J.V. directed research. F.P. and J.V. wrote the manuscript.

## Acknowledgements

We are very grateful to Anke Sparmann for her helpful comments on and editing of the manuscript. Further, we would like to thank Anna Nöhren and Esther Hauschild for their excellent technical assistance. We also would like to thank Svetlana Ðurica-Mitić, Kotaro Chihara and Gianluca Matera for their helpful comments and discussion. We thank the Vogel Stiftung Dr. Eckernkamp for supporting F.P. and V.C. with a Dr. Eckernkamp Fellowship. This work was supported by funds to J.V. from a DFG Gottfried Wilhelm Leibniz Award (DFG Vo875-18) and the Bavarian bayresq.net project Rbiotics.

---

**Figure S1:**
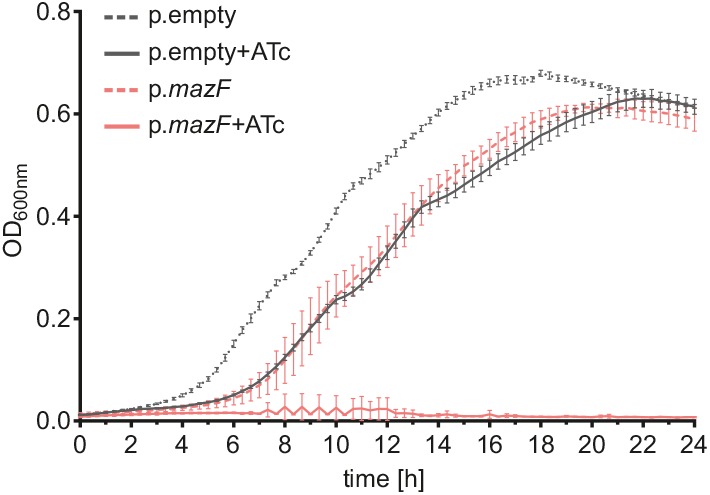
Testing of *mazF* toxicity during growth in the presence of selection pressure. Growth curve for *F. nucleatum* carrying either p. empty or p.*mazF* in the presence or absence of 100 ng ml^-1^ ATc. Thiamphenicol was added to select for plasmid maintenance. Shown is the optical density (OD_600nm_) over time (h, hours) for three biological replicates and the standard deviation.

**Figure S2:**
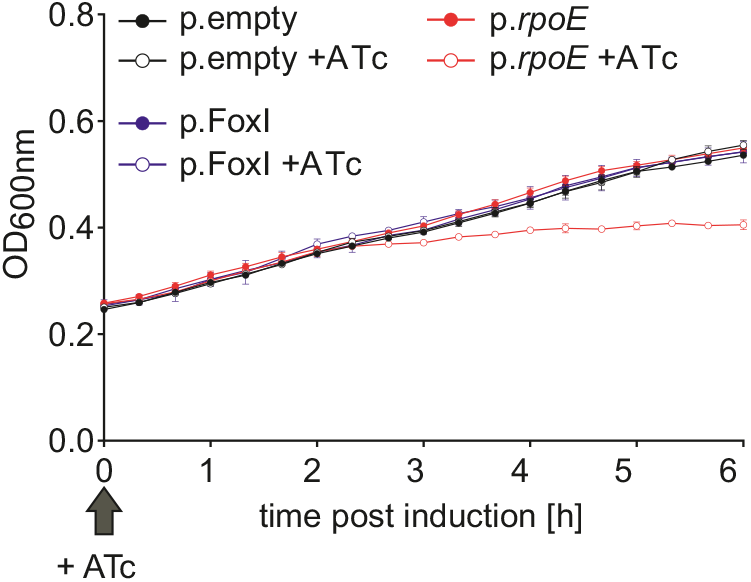
Impact of *rpoE* or FoxI inducible expression on growth. Growth curve for *F. nucleatum* carrying either p. empty, p.*rpoE* or p.FoxI in the presence or absence of 100 ng ml^-1^ ATc. The time point of ATc addition is indicated. Shown is the optical density (OD_600nm_) over time (h, hours) for three biological replicates and the standard deviation.

**Figure S3:**
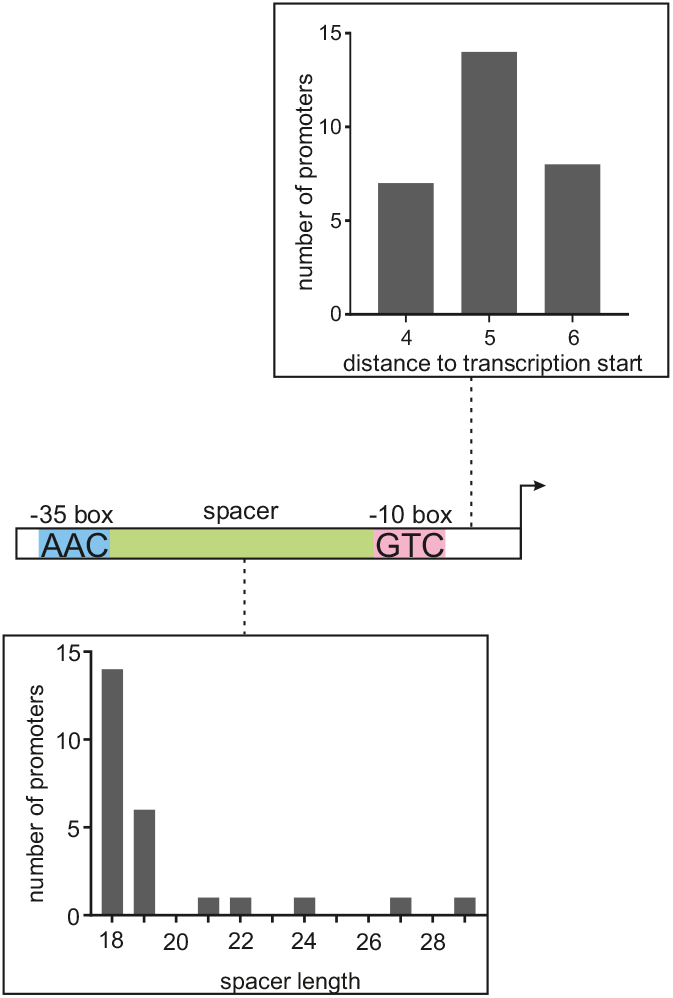
Analysis of spacing for the *rpoE* promoter motif. The bar charts display the distance between the -10 and -35 boxes (bottom) as well as from the -10 box to the TSS (top). Numbers of associated promotors are plotted on the Y-axis.

**Figure S4:**
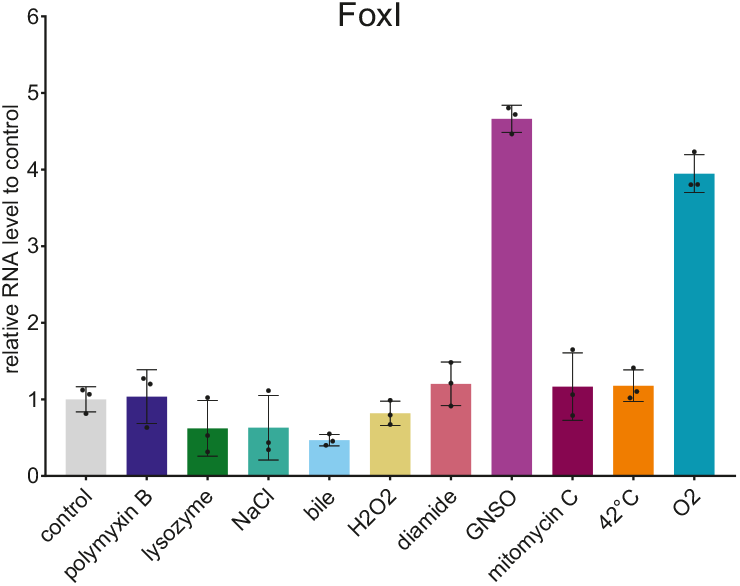
Quantification of FoxI levels upon exposure to different stress conditions. FoxI signal detected via northern blot was quantified for bacteria exposed to the indicated stress conditions. The average of three biological replicates relative to that of the control is displayed together with the standard deviation.

**Figure S5:**
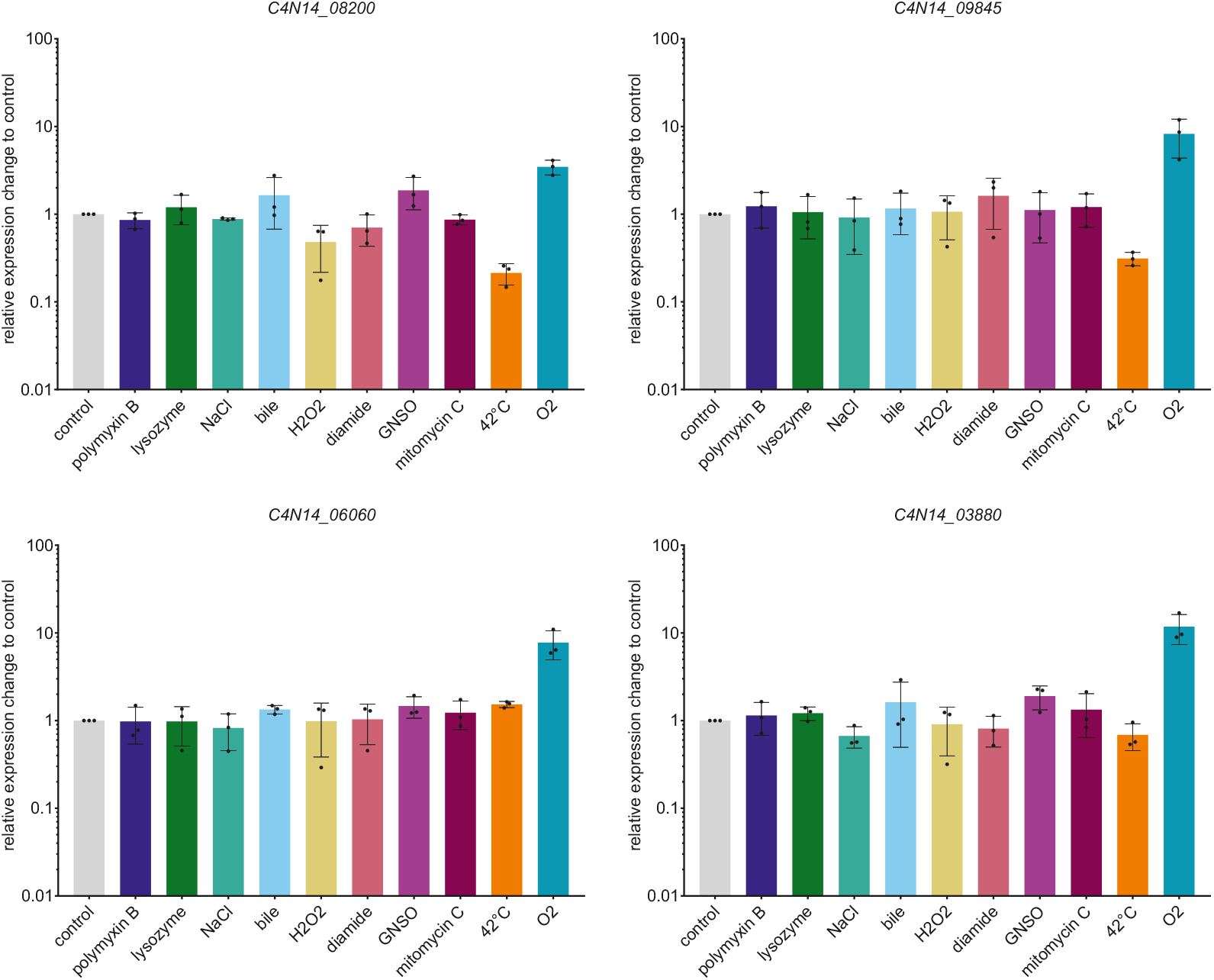
RT-qPCR analysis for additional genes upon exposure to different stress conditions. RT-qPCR analysis for mRNA of the indicated genes after exposing *F. nucleatum* to indicated stress conditions for 60 min. Data are normalized to the control and plotted as the average of three biological replicates with the standard deviation.

**Figure S6:**
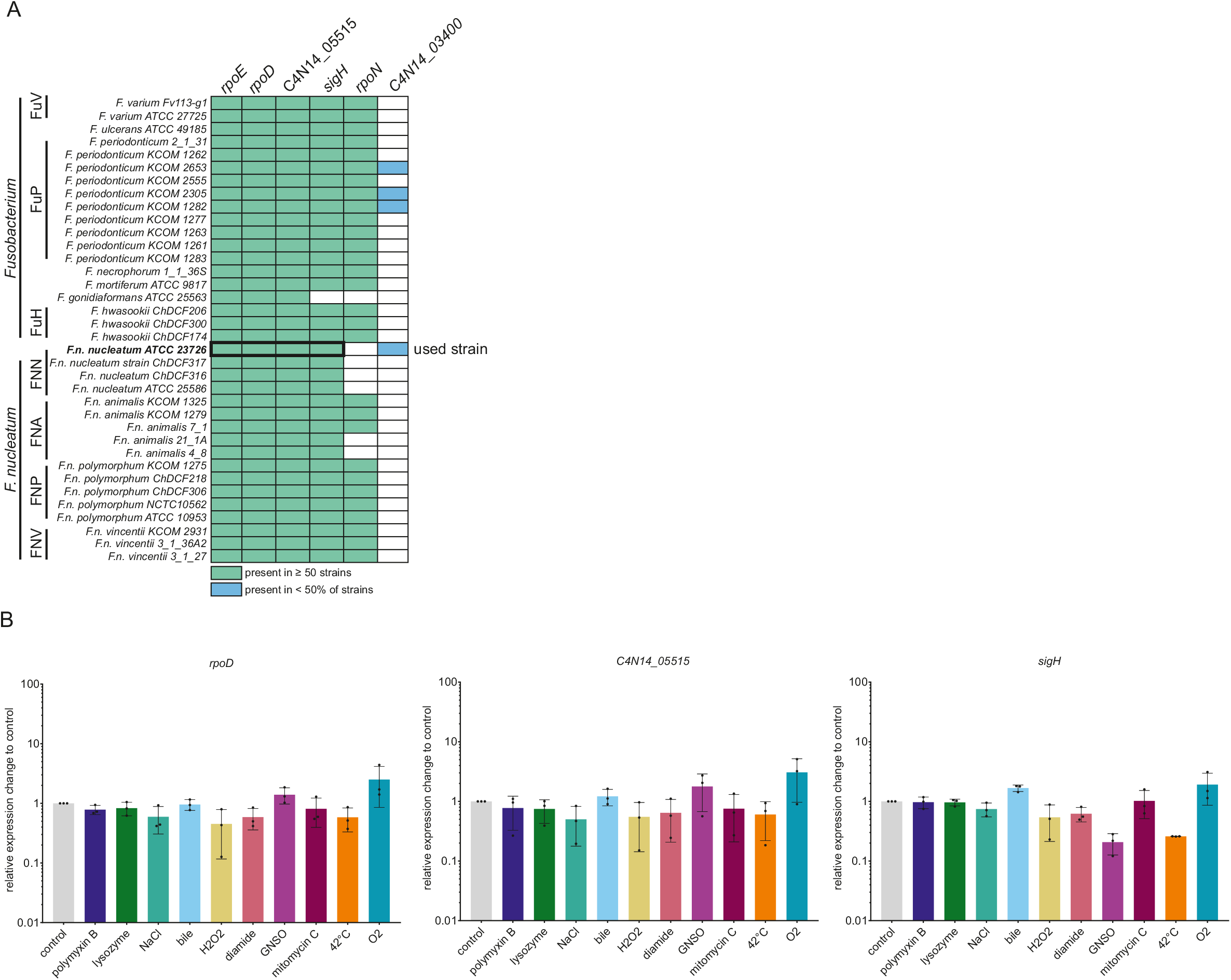
Response of sigma factors to the different stress conditions. (A) Schematic representation of the presence or absence of annotated sigma factors in species of *F. nucleatum* and members of the *Fusobacterium* genus. White fields mark the absence of the sigma factor. The black box indicates all conserved sigma factors investigated in this study.(B) RT-qPCR analysis for mRNA of the indicated genes after exposing *F. nucleatum* to indicated stress conditions for 60 min. Data are normalized to the control and plotted as the average of three biological replicates with the standard deviation.

**Figure S7:**
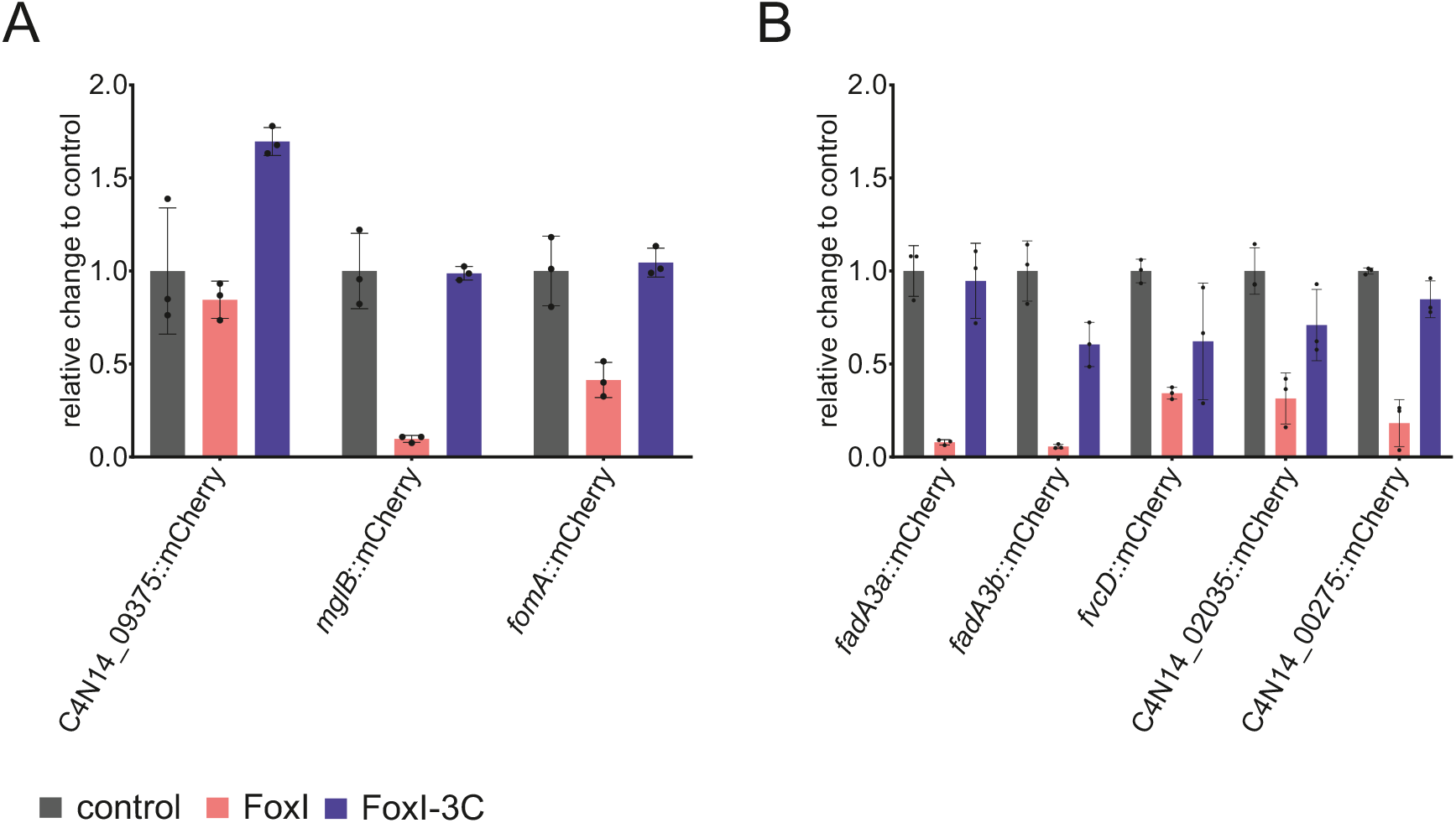
Western blot quantification for different translational reporters. Quantification of western blots probed for mCherry expressed from the translational reporter plasmids in the presence of FoxI or FoxI-3C analyzed in Figure 5 for (A) and Figure 6 for (B). 5’UTRs of the indicated genes were analyzed. The average of three biological replicates relative to that of the control is displayed together with the standard deviation.

**Figure S8:**
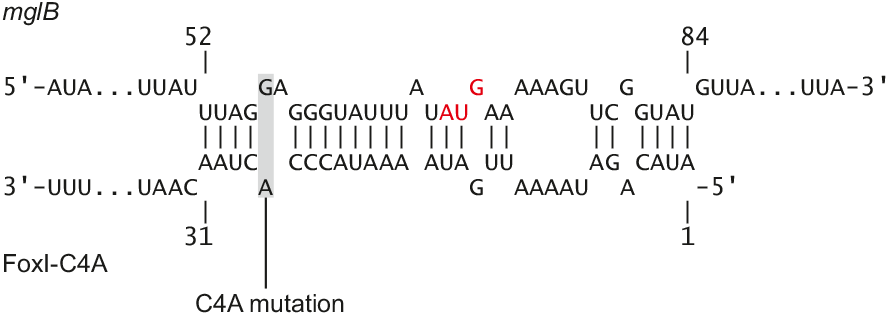
IntaRNA target prediction between *mglB* mRNA and FoxI-C4A. Schematic representation of the IntaRNA base-pairing prediction between the *mglB* mRNA and the seed region mutant FoxI-C4A. The site of mutation on the sRNA (grey) and the start codon of the mRNA (red) are indicated.

